# The Soybean Cyst Nematode Effector Cysteine Protease 1 (CPR1) Targets a Mitochondrial Soybean Branched-Chain Amino Acid Aminotransferase (GmBCAT1) for Degradation

**DOI:** 10.1101/2024.07.01.601533

**Authors:** Alexandra Margets, Jessica Foster, Anil Kumar, Tom R. Maier, Rick Masonbrink, Joffrey Mejias, Thomas J. Baum, Roger W. Innes

## Abstract

The soybean cyst nematode (SCN; *Heterodera glycines*) facilitates infection by secreting a repertoire of effector proteins into host cells to establish a permanent feeding site composed of a syncytium of root cells. Among the diverse proteins secreted by the nematode, we were specifically interested in identifying proteases to pursue our goal of engineering decoy substrates that elicit an immune response when cleaved by an SCN protease. We identified a cysteine protease that we named Cysteine Protease 1 (CPR1), which was predicted to be a secreted effector based on transcriptomic data obtained from SCN esophageal gland cells, presence of a signal peptide, and lack of transmembrane domains. CPR1 is conserved in all isolates of SCN sequenced to date, suggesting it is critical for virulence. Transient expression of CPR1 in *Nicotiana benthamiana* leaves suppressed cell death induced by a constitutively active nucleotide binding leucine-rich repeat protein, RPS5, indicating that CPR1 inhibits effector-triggered immunity. CPR1 localizes in part to the mitochondria when expressed *in planta*. Proximity-based labeling in transgenic soybean roots, co-immunoprecipitation, and cleavage assays identified a branched-chain amino acid aminotransferase from soybean (GmBCAT1) as a substrate of CPR1. Silencing of the *CPR1* transcript in the nematode reduced penetration frequency in soybean roots while the expression of *CPR1* in soybean roots enhanced susceptibility. Our data demonstrates that CPR1 is a conserved effector protease with a direct target in soybean roots, highlighting it as a promising candidate for decoy engineering.

## Introduction

The Soybean Cyst Nematode (SCN; *Heterodera glycines*) is an obligate endoparasite that is responsible for yield losses exceeding $1 billion annually in North America (Arjoune et al. 2022). Current methods to mitigate economic losses include crop rotations with non-host plants and the use of resistant soybean varieties. However, the overuse of resistant soybean varieties, which are primarily derived from a single source of resistance (PI 88788), have caused shifts in SCN populations (Niblack et al. 2008; Allen et al. 2017), diminishing the efficacy of these varieties (Mitchum et al. 2007; McCarville et al. 2017). Therefore, new methods of resistance are critical to ensure soybean yields long term. Development of durable SCN prevention strategies requires a deeper understanding of SCN virulence mechanisms.

Arguably, the establishment of a permanent feeding site, known as the syncytium, is the most important characteristic of SCN infection as it facilitates the long-term relationship between the plant and nematode. To form syncytia, cyst nematodes induce large-scale morphological changes in host plant cells, including loss of the central vacuole, expansion of the cytoplasm, digestion of cell walls, and protoplast fusion of neighboring cells (Jones 2008). The syncytium serves as a nutrient source providing sugars, starches, fatty acids, amino acids, and B vitamins necessary for nematode development and reproduction (Lin and Siddique 2024). Without a viable syncytium, cyst nematodes are unable to complete their life cycle.

To reprogram host cells and evade the plant immune system (Jones and Dangl 2006; Sato et al. 2019), cyst nematodes secrete a diverse repertoire of effector proteins. Effectors are broadly defined as proteins secreted by a pathogen during infection to alter the host cell’s structure and/or function (Hogenhout et al. 2009). In cyst nematodes, effectors are synthesized primarily in the esophageal gland cells (two sub-ventral and one dorsal) of the nematode and then secreted through a hollow mouth spear called the stylet. The sub-ventral gland cells are more prominent during early stages of infection and involved in the secretion of cell-wall degrading enzymes during root penetration (Pellegrin et al. 2024), whereas the dorsal gland cells play a pivotal role during sedentary stages and are thought to facilitate suppression of plant immune responses and initiating and maintaining the syncytium (Davis et al. 2004).

Nematologists have employed various approaches to identify effectors, which are reviewed in (Davis et al. 2004; Vieira and Gleason 2019). The availability of an annotated SCN genome (Masonbrink et al. 2019b) and the ability to isolate and purify nematode gland cells for transcriptomic analysis (Maier et al. 2013) have both facilitated identification of novel SCN effectors. Using these resources, we specifically aimed to identify SCN proteases that are secreted during infection and function inside plant cells, as our overarching goal is to engineer a plant-produced decoy substrate that, when cleaved by an SCN protease effector, elicits a plant immune response.

This decoy engineering approach is based on our discovery that cleavage of the *Arabidopsis* AvrPphB SUSCEPTIBLE 1 (PBS1) protein by the *Pseudomonas syringae* effector protease AvrPphB activates the nucleotide binding leucine-rich repeat (NLR) disease resistance protein RESISTANCE TO PSEUDOMONAS 5 (RPS5). Upon activation, RPS5 triggers a localized cell death known as the hypersensitive response (HR). We have previously shown that the AvrPphB cleavage sequence within the activation loop of PBS1 can be substituted with cleavage sequences specific to proteases secreted by other pathogens (Kim et al. 2016; Helm et al. 2019; Pottinger et al. 2020). The key discoveries enabling development of the PBS1 decoy technology are reviewed in (Pottinger and Innes 2020). Notably, PBS1 is highly conserved across flowering plants, as is recognition of AvrPphB (Carter et al. 2019), which has enabled development of similar decoy systems in crop species such as soybean (Helm et al. 2019; Pottinger et al. 2020). Because SCN relies on living cells to form a syncytium, we expected the PBS1 decoy system, which induces cell death upon activation, would be effective in conferring resistance to SCN. As a first step toward development of a PBS1 decoy effective against SCN, we needed to identify SCN effector proteases that contribute to virulence. Such proteases have been described in bacterial (Figaj et al. 2019), fungal (Chandrasekaran et al. 2016), viral (Rodamilans et al. 2018), and nematode (Antonino de Souza Júnior et al. 2014) plant pathogens. However, little is known about effector proteases in cyst nematodes specifically.

In addition to identifying candidate SCN effector proteases, we also aimed to identify the targets of these proteases inside soybean cells. Identification of such targets will provide insight into the plant processes that SCN modifies to promote virulence and will also aid in the identification of preferred cleavage sequences for each protease, which will be required for future decoy engineering.

Here we describe identification of an SCN effector protease, which we have named <u>C</u>ysteine <u>Pr</u>otease 1 (CPR1; Hetgly22189). Gland transcriptome analysis showed that CPR1 is expressed in parasitic-stage gland cells. Transient expression in *Nicotiana benthamiana* showed that CPR1 localizes, in part, to mitochondria and can suppress cell death triggered by RPS5. Proximity-based labeling experiments in transgenic soybean roots identified a soybean branched-chain amino acid aminotransferase (GmBCAT1; Glyma.06G050100) as a putative target, which was further confirmed through co-immunoprecipitation. Most significantly, co-expression of CPR1 with GmBCAT1 showed that GmBCAT1 is degraded by CPR1 in a protease-dependent manner. Silencing of *CPR1* in SCN reduced nematode penetration frequency, while expression of CPR1 in transgenic soybean roots enhanced susceptibility. These results show that CPR1 contributes to SCN virulence and represents an ideal candidate for development of a decoy substrate.

## RESULTS

### Identificatiown of CPR1

To identify candidate SCN effector proteases, we searched SCN esophageal gland cell transcriptomes for transcripts annotated as proteases. We further confirmed the presence of the protease domains using MEROPS peptidase and NCBI domain prediction databases. We selected those predicted to harbor signal peptides (SignalP5.0) and lack transmembrane domains (TMHMM 2.0). To further prioritize our list of proteases, we referred to previously generated RNA-seq data comparing esophageal gland cell expression to whole worm samples (Maier et al. 2021). Since SCN effectors are synthesized in gland cells (Gao et al. 2001; Noon et al. 2015), we selected proteases that were upregulated in glands compared to the whole worm. In this pursuit, we were especially interested in identifying and characterizing cysteine proteases, due to their previous success in PBS1 decoy engineering (Kim et al. 2016; Helm et al. 2019; Pottinger et al. 2020).

Of the proteases in the SCN gland cell library, CPR1 was the top priority candidate as it was annotated as a cysteine protease and highly conserved in the SCN genomes available (early release data available on SCNBase) (Masonbrink et al. 2019a). CPR1 is 565 amino acids long and predicted to belong to the C1 Peptidase superfamily (Fig. 1A). The protease harbors a cathepsin propeptide inhibitor domain (I29), which is known to form an alpha helical structure that blocks its own substrate binding site *in cis* to prevent it from binding to substrates. The catalytic cysteine residue was identified as cysteine 323 based on alignment to similar proteases that were identified in the MEROPS peptidase database. SignalP5.0 identified a signal peptide spanning from amino acids 1-16. TMTHH predicted that the protease lacked transmembrane domains (Fig. 1A).

**Fig. 1.**
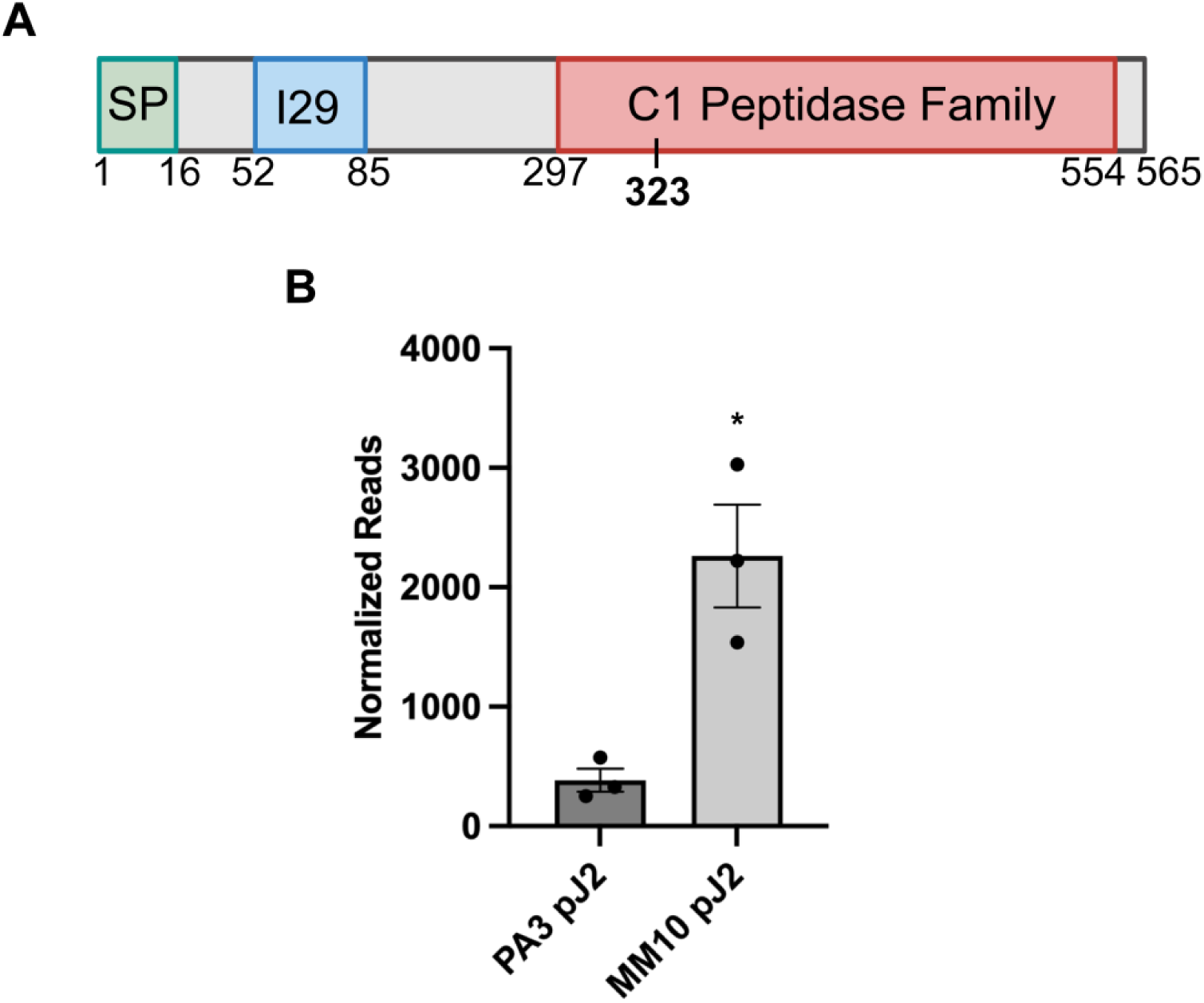
CPR1 is a gland synthesized cysteine protease. **A,** Schematic of the CPR1 protein. SP= signal peptide, I29= I29 inhibitor domain, C1 Peptidase= protease domain. Numbers represent amino acid position. Amino acid 323 (bold) is the catalytic cysteine residue. **B,** Normalized read counts of *CPR1* in glands of PA3 and MM10 populations at the parasitic J2 (pJ2) life stage. The data represent mean values ±SEM of three biological replicates for each population. Statistical analysis employed a two-tailed unpaired t-test, *P*<0.05 (*).

Analysis of gland cell libraries from two different SCN populations, PA3 and MM10, prepared from parasitic J2 (pJ2) nematodes, revealed that CPR1 transcripts are more abundant in glands of MM10 pJ2 worms (Fig. 1B), which was determined using normalized read counts from three biological replicates. This is noteworthy as SCN population MM10 is defined as an HG type 1-7 (Maier et al. 2021), meaning this population is capable of reproducing on all seven resistant soybean indicator lines (Niblack et al. 2003). SCN population PA3 is defined as an HG type 0, meaning this population is incapable of reproducing on all seven resistant soybean indicator lines. The elevated expression of CPR1 in the MM10 population suggests that CPR1 may aid in overcoming resistance.

### CPR1 suppresses RPS5-mediated cell death in *N. benthamiana* leaves

A common role of effectors is to suppress plant immune responses, including both pattern-triggered immunity (PTI) and effector-triggered immunity (ETI). To determine whether CPR1 can suppress ETI-dependent HR, we used an auto-active RPS5 mutant (RPS5^D266E^) that elicits a localized cell death response when overexpressed in *N. benthamiana* without an effector present (Qi et al. 2012). To perform these assays, we first co-expressed RPS5^D266E^ with either CPR1 (lacking its signal peptide) or an empty vector (ev) control. At 24 h post gene expression induction, the area of the leaf infiltrated with RPS5^D266E^ and CPR1 lacked cell death while the area infiltrated with RPS5^D266E^ and the ev control exhibited complete tissue collapse. We confirmed that the area infiltrated with the protease contained living cells using ultraviolet light as the chlorophyll auto-fluoresces red in living cells (Fig. 2A). This result showed that in the presence of CPR1, ETI-dependent HR was suppressed and that CPR1 can function inside of *N. benthamiana* cells. While screening CPR1 for ETI suppression activity, we screened a second SCN protease in parallel (20453) that was unable to suppress RPS5-mediated HR among all biological replicates (Supplementary Fig. S1A). 20453 served as a second negative control for this assay. To test whether CPR1’s HR suppression activity required protease function, we mutagenized the catalytic cysteine residue of CPR1 to a serine at amino acid position 323. Co-expression of RPS5^D266E^ with the inactive protease (CPR1^C323S^) resulted in complete tissue collapse (Fig. 2B), indicating that the protease activity of CPR1 is required for HR suppression.

**Fig. 2.**
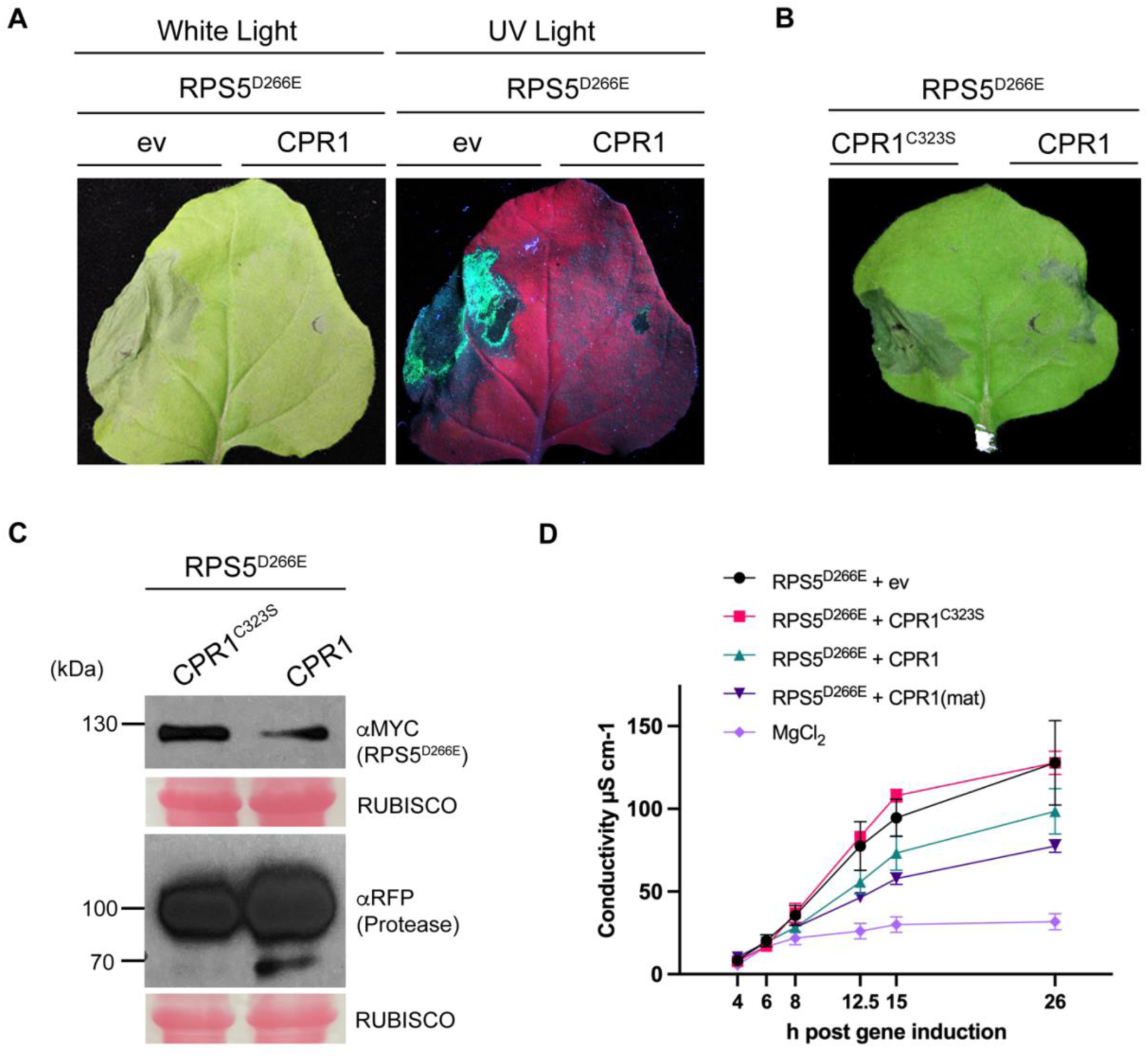
CPR1 suppresses ETI-dependent cell death. **A**, Cell death induced by auto active RPS5 (RPS5^D266E^:5xMYC) is suppressed when co-expressed with CPR1:mCherry. The indicated constructs were transiently expressed in *N. benthamiana* using dexamethasone-inducible promoters. Leaf images are representative of 10 leaves. 10/10 leaves co-expressing RPS5^D266E^ and empty vector (ev), and 0/10 leaves co-expressing RPS5^D266E^ and CPR1 displayed tissue collapse. Images were taken under white light and ultraviolet (UV) light. **B,** RPS5^D266E^ mediated cell death is not suppressed by the protease inactive variant (CPR1^C323S^). 8/8 leaves co-expressing RPS5^D266E^ and CPR1^C323S^ and 2/8 leaves co-expressing RPS5^D266E^ and CPR1 displayed tissue collapse. Images in **A** and **B** were taken 24 h post gene expression induction. **C,** Immunoblot showing expression of the RPS5^D266E^ and CPR1^C323S^ or CPR1. RUBISCO was used as a loading control. **D,** Electrolyte leakage assay of *N. benthamiana* leaves co-infiltrated with RPS5^D266E^ and ev, CPR1, CPR1^C323S^, or CPR1(mat). MgCl_2_ treated leaves were used as a buffer control. Readings were taken from 4 individual leaf samples, containing 3 leaf discs each from different areas of the leaf avoiding any mechanical damage near injection sites. Error bars represent ±SEM. Electrolyte leakage assay was repeated twice with consistent results.

Suppression of the RPS5-induced HR by CPR1 could potentially be caused by interference with RPS5 accumulation, either via destabilization or inhibition of transient transformation. To rule out these explanations, we assessed RPS5^D266E^ accumulation using an immunoblot. Neither wild-type CPR1 nor CPR1^C323S^ inhibited RPS5^D266E^ accumulation (Fig. 2C). Since the expression level of full-length RPS5^D266E^ was similar in both the inactive and active protease samples and because RPS5 is not conserved in soybean, RPS5 is unlikely to be a direct target of CPR1.

Immunoblot analyses of the above samples also revealed a putative self-cleavage product in the wild type CPR1 protein at approximately 70 kDa that was missing from the CPR1^C323S^ samples (Fig. 2C). Based on its size, such self-processing would remove the N-terminal inhibitor domain of CPR1, and thus would be expected to activate it. We therefore generated a “mature” variant (CPR1(mat)) lacking the inhibitor domain to mimic the processed version of the protease.

To further assess whether protease activity and proteolytic processing influenced the cell death suppression observed, we transiently co-expressed RPS5^D266E^ with ev, CPR1, CPR1^C323S^, or CPR1(mat) in *N. benthamiana* leaves. Three leaf discs from four different leaves per condition were harvested approximately 3.5 h post gene expression induction and an electrolyte leakage assay was performed to measure cell death via conductivity. Readings were taken at 4, 6, 8, 12.5, 15, and 26 h post gene expression induction. As expected, the samples containing RPS5^D266E^ co-expressed with either ev or CPR1^C323S^ resulted in the highest levels of cell death. When RPS5^D266E^ was co-expressed with CPR1 or CPR1(mat), reduced cell death was observed. Notably, the samples expressing CPR1(mat) resulted in the lowest levels of cell death, except for the MgCl_2_ control (Fig. 2D). To further support the conclusion that the lack of cell death was not due to lack of expression of RPS5^D266E^, representative leaf tissue was harvested from the electrolyte leakage assay and subjected to immunoblot analysis. All samples were found to be expressing the protease variants and RPS5^D266E^ at similar levels (Supplementary Fig. S1B). Together, these results show that CPR1 is capable of suppressing RPS5-mediated HR in a protease-dependent manner. Furthermore, removal of the inhibitor domain of CPR1 enhances its cell death suppression activity.

### CPR1 localizes to the cell periphery in puncta-like structures in *N. benthamiana* leaves

Because our long-term goal is to generate a PBS1 decoy protein that can be cleaved by an SCN protease, we assessed whether CPR1 co-localizes with the soybean PBS1 protein (GmPBS1a), as partial co-localization is needed to facilitate cleavage of PBS1 (Qi et al. 2014; Pottinger et al. 2020). When GmPBS1a:mCherry was transiently co-expressed with CPR1(mat):sYFP in *N. benthamiana* leaves, the CPR1:sYFP fluorescence overlapped with the GmPBS1a:mCherry fluorescence at the cell periphery, but we also observed numerous puncta-like structures (Fig. 3A) reminiscent of mitochondria. To assess whether CPR1 might be partially localizing to mitochondria, we stained *N. benthamiana* leaves expressing CPR1(mat):sYFP with MitoTracker Red. The CPR1 puncta co-localized with the mitochondrial marker (Fig. 3B), confirming that CPR1 partially localizes to mitochondria. Consistent with this observation, analysis of the CPR1(mat) sequence using iPSORT (Bannai et al. 2002) revealed a predicted mitochondrial localization signal. Although CPR1 appears to be partially localized to mitochondria, its presence at the cell periphery suggests that it should also have access to PBS1, which is targeted to the plasma membrane by acylation on its N-terminus (Qi et al. 2014).

**Fig. 3.**
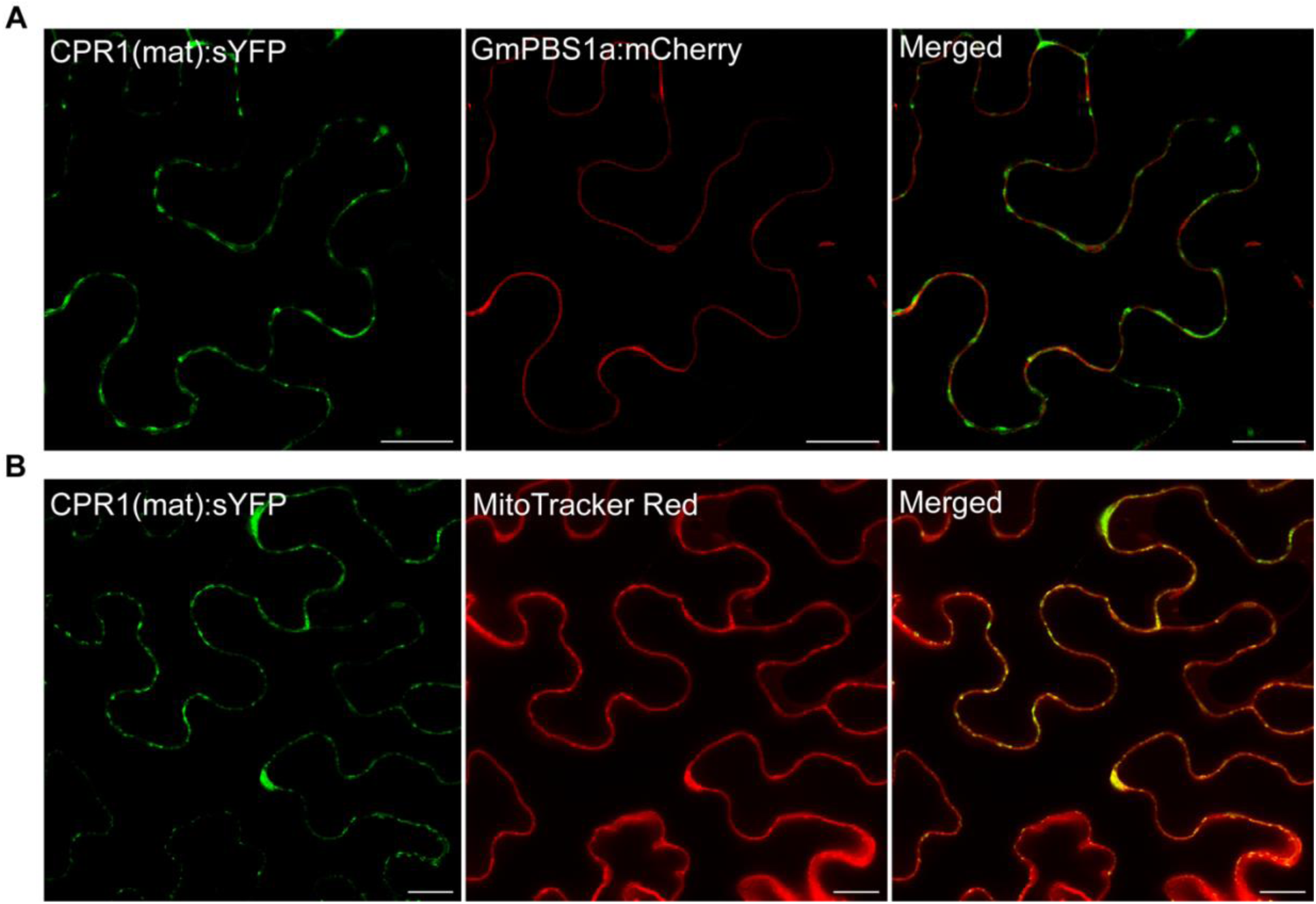
CPR1(mat) co-localizes with GmPBS1a at the cell periphery and aggregates in puncta-like structures in *N. benthamiana* leaves. **A,** Confocal images showing the expression and subcellular co-localization of GmPBS1a:mCherry and CPR1(mat):sYFP in *N. benthamiana* leaves. **B,** Confocal images showing the expression of CPR1(mat):sYFP and mitochondria stained with MitoTracker Red. Images in **A** and **B** were collected 5 to 8 h post induction of gene expression by dexamethasone application and 1 h post injection of MitoTracker Red. Scale bars = 20 μm.

### Identification of candidate substrates of CPR1 using biotin-mediated proximity labeling

To identify potential substrates of CPR1 *in planta*, we used a proximity-based biotin labeling system (miniTurbo; mT) (Zhang et al. 2020) that we optimized for use in transgenic soybean roots. Specifically, we generated composite soybean plants consisting of wild-type shoots and transgenic roots expressing CPR1 variants fused to a miniTurbo biotin ligase (CPR1:miniTurbo). To facilitate identification of transgenic roots, we included the *RUBY* reporter gene in our T-DNA construct (He et al. 2020). By 21-32 days post inoculation (dpi) of soybean hypocotyls with *Agrobacterium rhizogenes* K599 strains, RUBY-colored hairy roots expressing CPR1 (with and without protease activity and with and without the inhibitor domain), were readily observed (Supplementary Fig. S2). These composite plants were then treated with exogenous biotin, and the roots were harvested 4 h later. As a negative control, we generated composite soybean plants expressing YFP:miniTurbo. Biotinylated proteins were purified using streptavidin beads and prepared for mass spectrometry. A total of 277 soybean proteins were identified by mass-spectrometry analysis across all four of the CPR1:miniTurbo variant samples. Of these, we focused on proteins that were present in CPR1 datasets and absent in datasets of other SCN effectors being pursued for projects external to this study. Based upon our analysis, we selected 12 soybean proteins to test for direct interaction and cleavage (Supplementary Table S1). Ten out of twelve of these proteins were enriched in CPR1 datasets compared to the YFP:miniTurbo datasets. While we included different CPR1 variants with and without protease activity and/or the inhibitor domain, there were no significant enrichments between the CPR1 samples that influenced our selection of top priority candidates. Notably, several of the selected proteins have links to SCN infection and/or plant immunity from prior studies. Using soybean cDNA, we successfully cloned seven of the twelve proteins for further analysis.

### CPR1 interacts with GmBCAT1

To confirm interactions between CPR1 and the soybean proteins identified from the miniTurbo experiments, we first performed co-immunoprecipitation (co-IP) assays in *N. benthamiana* leaves. To stabilize the putative interactions, we used the mature variant of CPR1 lacking protease activity (CPR1(mat)^C323S^) with a C-terminal GFP tag as a bait protein. As a negative control, inactive AvrPphB (AvrPpB^C98S^), was used as the bait protein. Of the proteins identified in the miniTurbo datasets, we prioritized soybean branched-chain amino acid aminotransferase 1 (GmBCAT1) (Glyma.06G050100), as it was absent in the YFP:miniTurbo negative control and all other SCN effector datasets. We cloned GmBCAT1 from soybean root cDNA but left off the N-terminal 74 amino acids (GmBCAT1^Δ1-74^), which was predicted to contain a transmembrane domain that could potentially complicate co-IP analysis. GmBCAT1^Δ1-74^ was found to co-IP with CPR1(mat)^C323S^ but not with AvrPphB^C98S^ (Fig. 4A). In parallel to testing the interaction between GmBCAT1 and CPR1, we also tested a second putative soybean target that was annotated as a CMP/dCMP-type deaminase domain-containing protein (Glyma.09G080100), using the same bait proteins. Glyma.09G080100 did not co-IP with CPR1, further supporting the specificity of CPR1’s interaction with GmBCAT1 (Supplementary Fig. S3).

**Fig. 4.**
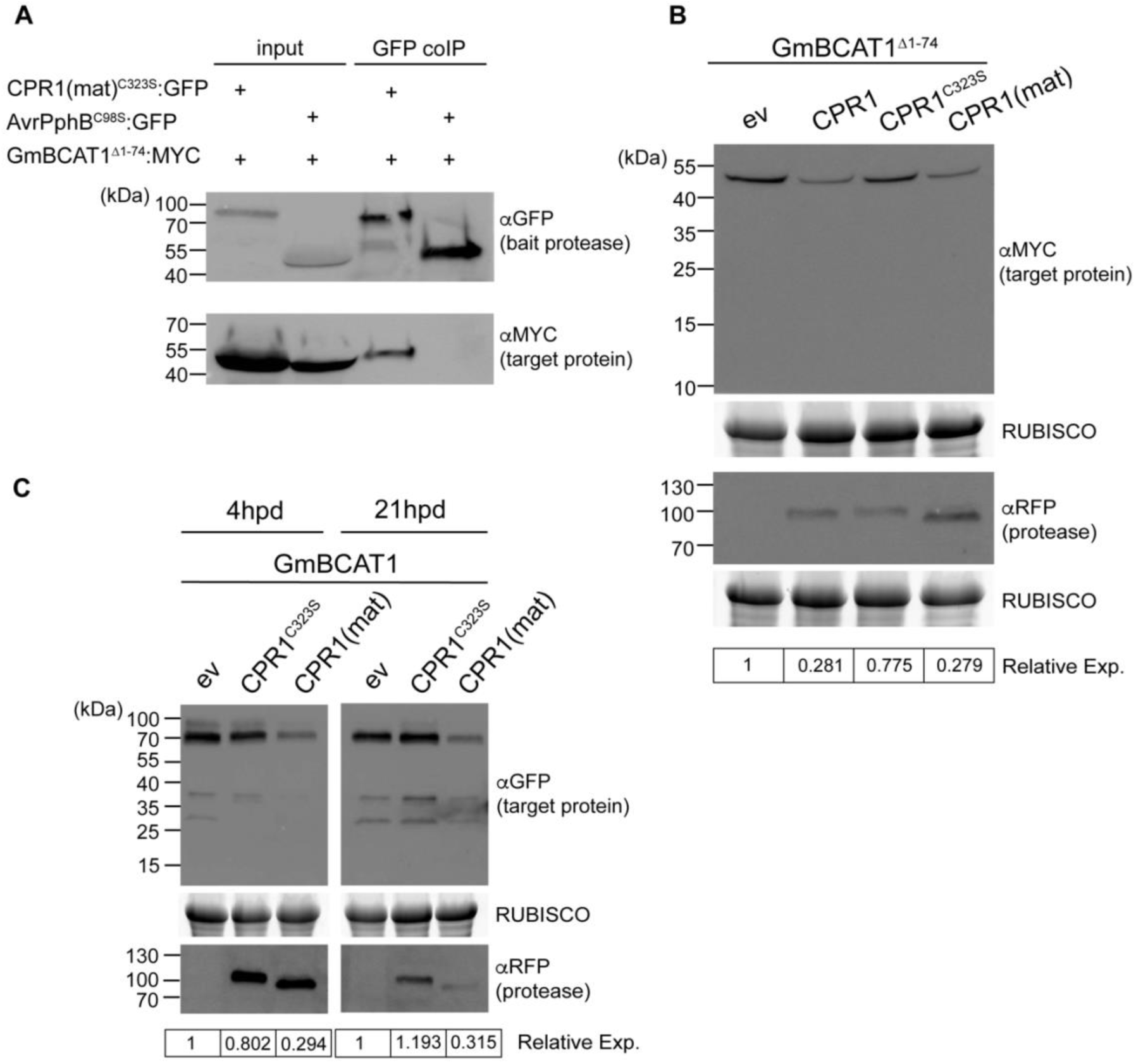
CPR1 targets a soybean branched-chain amino acid aminotransferase. **A,** GmBCAT1^Δ1-74^:4xMYC co-immunoprecipitates (co-IPs) with the protease-inactive CPR1(mat)^C323S^:GFP. The indicated proteins were transiently co-expressed in *N. benthamiana* and leaves harvested 21 h post dexamethasone application. GFP-tagged bait proteases were immunoprecipitated using GFP-Trap agarose beads. AvrPphB^C98S^ was used as a negative control. Co-IPs were repeated three times with consistent results. **B,** GmBCAT1^Δ1-74^ accumulation is reduced when co-expressed with active CPR1 variants. GmBCAT1^Δ1-74^ was co-expressed with empty vector (ev), active CPR1, protease inactive CPR1^C323S^, or CPR1(mat). **C,** GmBCAT1 (full-length minus signal peptide) accumulation is reduced when co-expressed with CPR1(mat) but not ev or CPR1^C323S^. Samples were harvested at 4 h and 21 h post dexamethasone application. In **B** and **C**, relative expression values of GmBCAT1 were normalized to the ev control and quantified using ImageJ. RUBISCO was used as a loading control. Cleavage tests were performed at least twice with consistent results. All constructs used were under control of dexamethasone inducible promoters.

To further characterize the interaction between CPR1 and GmBCAT1, we assessed whether GmBCAT1 protein could be proteolytically cleaved by CPR1. When GmBCAT1^Δ1-74^ was transiently co-expressed with CPR1 or CPR1(mat) in *N. benthamiana* leaves, a consistent reduction of GmBCAT1^Δ1-74^ protein accumulation was detected in comparison to the samples expressing the empty vector (ev) control or CPR1^C323S^ (Fig. 4B). We also transiently co-expressed the full-length version of GmBCAT1 with ev, CPR1, or CPR1(mat) in *N. benthamiana* leaves. Samples were harvested at 4 and 21 h post gene induction to assess whether the reduction of GmBCAT1 was maintained over multiple hours. As expected, accumulation of full-length GmBCAT1 was reduced in the presence of CPR1(mat) at 4 h post induction and maintained through the 21-h time-point (Fig. 4C). Notably, there was no detectable reduction in GmBCAT1^Δ1-74^ accumulation when expressed with the CPR1(mat)^C323S^ bait protein (Fig. 4A), further indicating that the protease activity facilitates GmBCAT1 degradation.

We used LOCALIZER (Sperschneider et al. 2017) to predict the subcellular localization of GmBCAT1. LOCALIZER returned no predicted targeting sequences for GmBCAT1. However, in a study that showed GmBCAT1 is upregulated upon drought stress (Do et al. 2022), iPSORT was used to identify a putative mitochondrial localization signal. We thus assessed whether GmBCAT1 localizes to mitochondria using MitoTracker Red and observed co-localization in puncta (Fig. 5A). The localization of CPR1 in mitochondria (Fig. 3B) strongly suggest this is the site in which targeting of GmBCAT1 occurs.

**Fig. 5.**
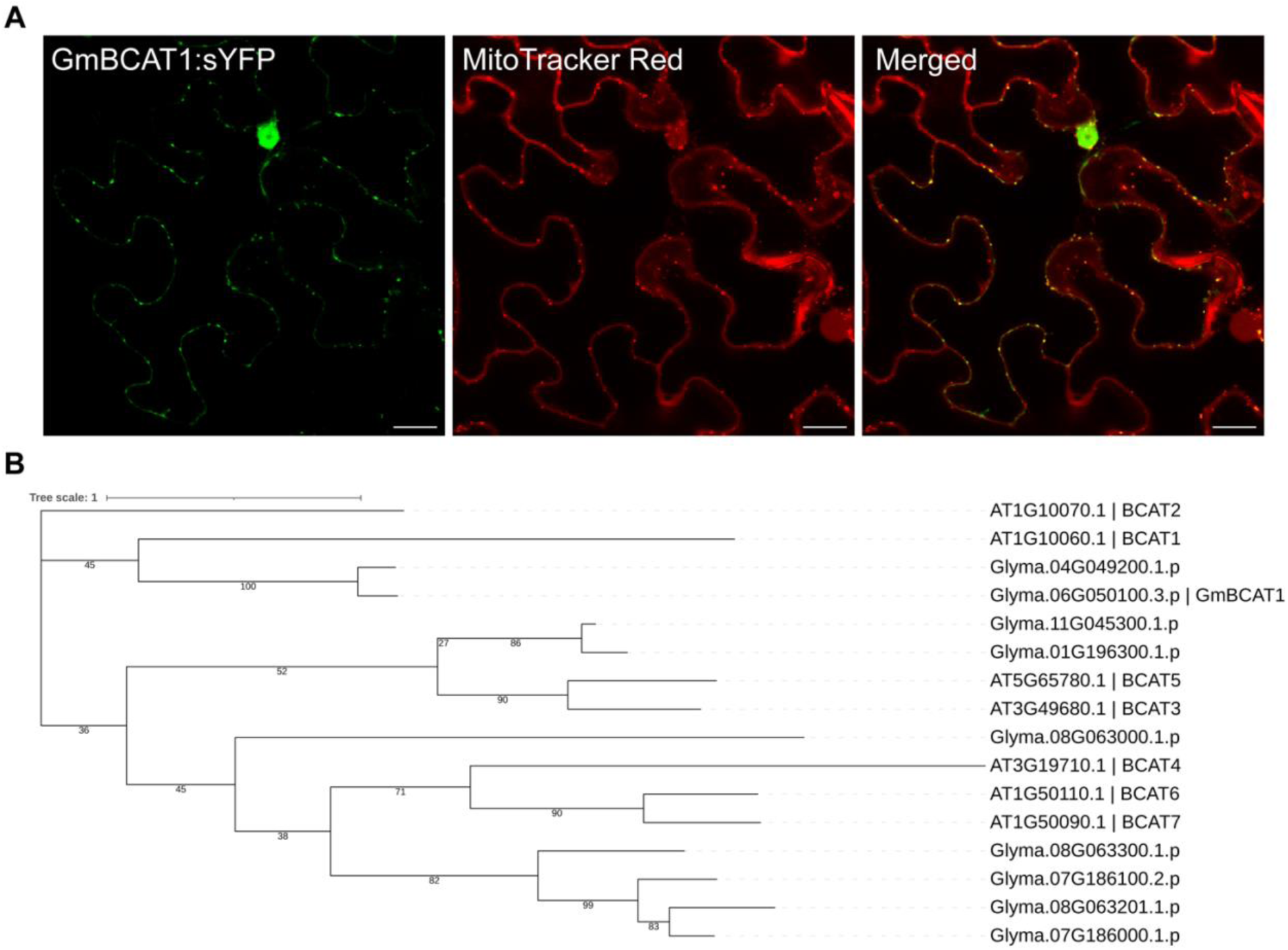
GmBCAT1 is a mitochondrial BCAT. **A,** Confocal images showing the subcellular co-localization of GmBCAT1:sYFP and mitochondria stained with Mitotracker Red approximately 5-6 hours post dexamethasone application and 1 hour post addition of Mitotracker Red. Scale bars = 20 μm **B,** Maximum Likelihood phylogenetic analysis of soybean and Arabidopsis BCAT proteins. Numbers at branch points represent percent total of 1000 bootstraps that support the indicated tree topology.

Since there is less known about soybean BCATs in comparison to those identified in *Arabidopsis*, we performed phylogenetic analysis to identify GmBCAT1 orthologs (Fig. 5B). A total of 9 soybean BCATs were identified, along with the 7 previously described *Arabidopsis* BCATs (Diebold et al. 2002). GmBCAT1 was determined to be most closely related to *Arabidopsis* BCAT1. Notably, Glyma.04G049200.1p is 95% identical GmBCAT1 at the amino acid level, and thus likely represents a homoeolog derived from the most recent genome duplication event in soybean.

### CPR1 contributes to SCN virulence

To assess whether CPR1 contributes to the virulence of SCN, we used RNA interference (RNAi) technology to silence *CPR1*. Designing a gene-specific probe is crucial for RNAi to avoid off-target effects, so we first conducted a BLASTn analysis of *CPR1* to identify similarities in the SCN genome (https://blast.scnbase.org/blastn). A gene-specific probe was then designed to specifically target *CPR1*. This probe template was then amplified using PCR resulting in a *CPR1*-specific amplicon. In brief, a dsRNA probe targeting a region common to all *CPR1* isoforms was synthesized using *in vitro* transcription. Since the optimal dsRNA concentration for efficient gene silencing was uncertain, we tested two concentrations: 2 mg/ml or 3 mg/ml of the *CPR1* dsRNA. After soaking, the nematodes were harvested, washed, and subjected to RT-qPCR to assess the effect of dsRNA soaking on the *CPR1* transcript. Soaking in dsRNA resulted in an approximately 85% reduction in *CPR1* transcripts for both concentrations of dsRNA tested (Fig. 6A).

**Fig. 6.**
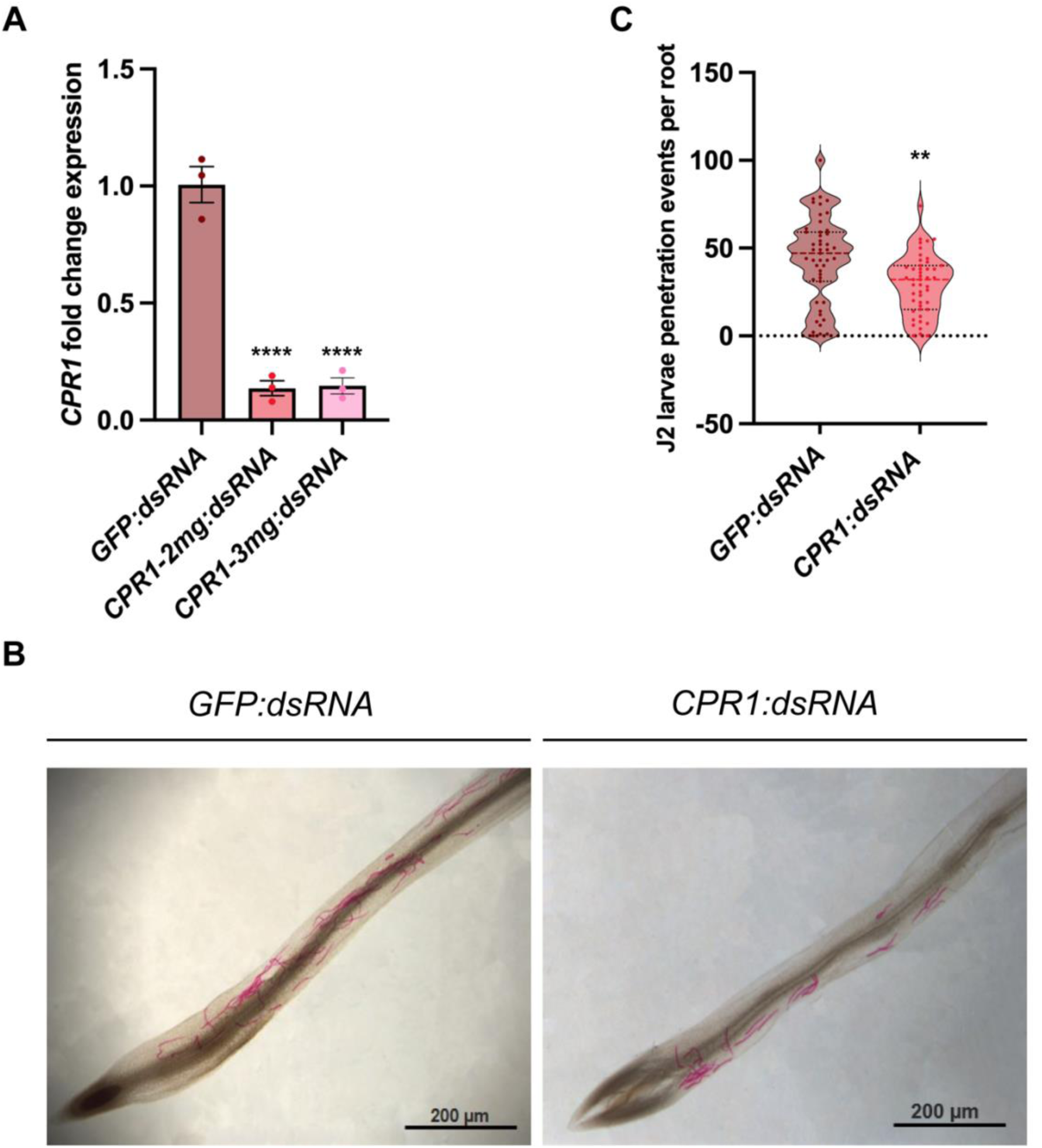
Knockdown of *CPR1* reduces *H. glycines* penetration of soybean roots. **A,** qRT-PCR analysis of *CPR1* transcript abundance in *H. glycines* J2s after soaking in *CPR1-dsRNA* (2 mg/ml and 3 mg/ml concentration) and *dsRNA-GFP* (negative control) at 24 h. Three biological and three technical replicates were included. This experiment was repeated three times with similar results. The *H. glycines* GAPDH gene (CA939315.1) was used for data normalization across samples. The data represent mean values ±SEM. Statistical analyses employed One-way ANOVA tests, with significance set as *P*<0.05. A Dunnett’s multiple comparison test showed *P* was significant at <0.0001 (****). **B,** Nematode penetration assay on soybean roots. J2 worms that had been soaked in dsRNA were inoculated onto root tips of 5-day old soybean seedlings placed in a pluronic gel (23% Pluronic F-127) and then incubated for 24 h. Roots were then stained with acid fuchsin to reveal the nematodes. **C,** *CPR1*-silenced nematodes display reduced penetration of soybean roots in comparison to controls (J2 worms treated with *dsRNA* targeting GFP). Statistical analysis employed an unpaired t-test. *P*<0.01 (**).

We next investigated the effect of *CPR1* gene silencing on nematode parasitism. We used *CPR1-*silenced nematodes to perform penetration assays by infecting soybean seedlings with approximately 120 *CPR1*-silenced or control J2 nematodes. After allowing the nematodes to infect soybean seedlings for 24 h, the nematodes inside the roots were stained with acid fuchsin (Bybd et al. 1983) and counted (Fig. 6B). Seedlings inoculated with *CPR1*-silenced J2 nematodes showed a significant decrease (33%, *P*=0.0011) in penetration ability compared to those treated with GFP dsRNA as a control (Fig. 6C). Together these results suggest that *CPR1* expression promotes early stages of infection.

To further assess CPR1’s contribution to virulence, we generated soybean composite plants (Fan et al. 2020) with transgenic roots that were expressing CPR1:mCherry+RUBY, CPR1^C323S^:mCherry+RUBY, or mCherry+RUBY. Approximately 2000 SCN TN10 eggs were inoculated on 23-day old plants. Nematodes were allowed to infect and develop for 28 days prior to harvesting. Cysts per gram of root tissue were counted for each biological replicate (Fig. 7A). Roots expressing CPR1 or CPR1^C323S^ resulted in a significant increase in susceptibility compared to roots expressing mCherry+RUBY alone (Fig. 7B), further indicating that expression of the protease, even without protease activity, enhances virulence.

**Fig. 7.**
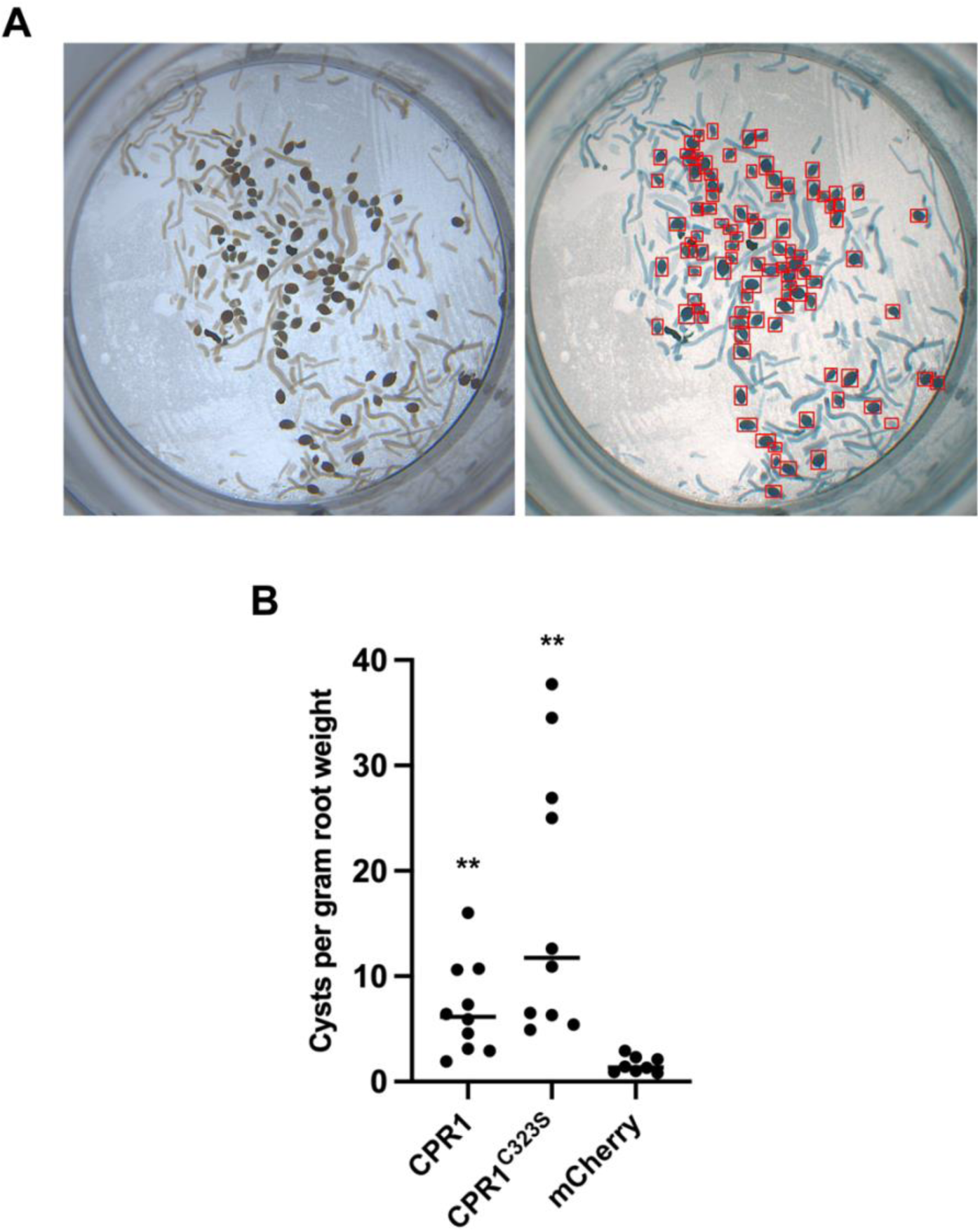
Expression of *H. glycines CPR1* in soybean roots enhances nematode susceptibility. Transgenic roots of individual composite plants growing in a sand:soil mixture were inoculated with 2,000 eggs per plant of SCN population TN10 and then plants grown for four additional weeks. Cysts were then collected and quantified as described in Materials and Methods. **A,** Representative images of cysts harvested and counted from *CPR1*-expressing roots using Nemacounter, which was designed in house at Iowa State University. Images show a mixed population of root debris and cysts (left) and the Nemacounter program detecting and counting the cysts specifically (right). **B,** Cysts per gram of soybean root weight were measured from roots expressing CPR1:mCherry+RUBY, CPR1^C323S^:mCherry+RUBY, and mCherry+RUBY. Roots were collected from 10, 10, and 8 individual composite soybean plants, respectively. Statistical analysis employed a Welch’s unpaired, t-test comparing roots expressing CPR1 or CPR1^C323S^ to control roots expressing mCherry. *P*<0.01 (**).

## DISCUSSION

Identifying targets of SCN effectors is valuable because it provides a snapshot into the biological processes that are manipulated by the nematode during infection. In addition, identification of CPR1 targets will enable determination of its preferred cleavage sequence, which could then be inserted into a PBS1 decoy. Prior to this work, yeast-two-hybrid assays have commonly been used to identify cyst nematode effector targets. We elected to use biotin ligase-based proximity labeling instead, as this approach enables detection of proteins that directly interact or come in close proximity to CPR1 inside soybean roots, where the protease normally functions during infection.

To express CPR1 in soybean roots, we used a composite plant system (Fan et al. 2020). Generation of transgenic roots using this approach requires just four to five weeks, which is much faster than the approximate 12 months required to generate whole transgenic soybean plants. In addition, this system does not require use of aseptic conditions, unlike more commonly used hairy root cultures (Noon et al. 2016), which are highly susceptible to contamination. The composite plant system, along with the MoClo Plant Tool Kits (Weber et al. 2011; Werner et al. 2012; Engler et al. 2014) allowed us to express CPR1 variants with a C-terminal biotin ligase (miniTurbo) and the visible marker RUBY on a single T-DNA. The miniTurbo system, used to identify GmBCAT1 as a target for CPR1, can identify transient interactions and resolve protein networks and complexes (Branon et al. 2018; Zhang et al. 2020). As we expected protease-substrate interactions to be transient, we viewed miniTurbo as a promising approach for identification of CPR1 target(s).

Proximity labeling enabled us to identify GmBCAT1 as a putative target of CPR1. The mature-inactive protease variant, CPR1(mat)^C323S^, was shown to interact with GmBCAT^Δ1-74^ (Fig. 4A). When co-expressed with CPR1 or CPR1(mat), GmBCAT^Δ1-74^ exhibited a significant reduction in accumulation in comparison to the empty vector control or a sample expressing the inactive protease, CPR1^C323S^ (Fig. 4B). While we did not observe a GmBCAT1 cleavage product when co-expressed with active CPR1 variants (Fig. 4B and C), a time-course cleavage assay showed that targeting of GmBCAT1 is rapid and sustained over a 21-h period post gene expression induction (Fig. 4C). Additionally, there was no reduction in GmBCAT^Δ1-74^ accumulation when expressed with the CPR1(mat)^C323S^ bait protein (Fig. 4A), further indicating that GmBCAT1 is targeted for degradation via CPR1 protease activity. The failure to observe a cleavage product suggests that such products are rapidly degraded.

BCATs are enzymes that catalyze the last step of leucine, isoleucine, and valine (branched-chain amino acids; BCAA) biosynthesis or the first step of their catabolism. In *Arabidopsis*, five of the six functional BCATs identified have been shown to localize to either the chloroplast, cytosol, or mitochondria (Diebold et al. 2002). GmBCAT1 has previously been shown to be upregulated upon drought stress and has been proposed to regulate induction of autophagy during stress responses (Do et al. 2022). Similarly, BCATs from rice have been shown to be induced upon drought stress (Shim et al. 2023), while a mitochondrial BCAT from wheat (*TaBCAT1)* was found to be a positive regulator of susceptibility to the wheat rust fungus (Corredor-Moreno et al. 2021). Our phylogenetic analysis revealed GmBCAT1 to be orthologous to *Arabidopsis* BCAT1 (Fig. 5B), which localizes to mitochondria and contributes to BCAA degradation (Schuster and Binder 2005; Binder 2010). If GmBCAT1 also participates in BCAA degradation, CPR1 may be targeting GmBCAT1 to maintain high levels of BCAAs in cysts and thus enhance their availability. Notably, BCAA levels in syncytia induced by *Heterodera schachtii* on *Arabidopsis* roots are elevated compared to uninfected roots (Anwar et al. 2016). In the same study, enzymes involved in BCAA biosynthesis, including *Arabidopsis* BCAT3 (Knill et al. 2008), were found to be upregulated in syncytia compared to uninfected roots. This increase in biosynthetic BCATs combined with proteolytic removal of catabolic BCAT1 by CPR1 could account for the elevated levels of BCAAs observed in syncytia.

To further assess CPR1’s contribution to virulence, we used RNAi to silence *CPR1* in J2-stage nematodes and then assessed the impact of *CPR1* silencing on nematode penetration frequency. Unlike RNAi protocols used for the free-living nematode *C. elegans,* techniques to silence plant-parasitic nematode effectors are limited. In the case of SCN and other plant-parasitic nematodes, techniques to perform RNAi are more involved and generally employ host-induced gene silencing or, alternatively, soaking of nematodes in dsRNA. A previous study successfully knocked-down proteases in the root knot nematode *Meloidogyne incognita* using plant-derived dsRNA, which substantially reduced the number of eggs per gram of root (Antonino de Souza Júnior et al. 2014). We used the dsRNA soaking approach to silence *CPR1.* The reduced number of J2 larvae penetration events in the soybean roots infected with *CPR1*-silenced nematodes compared to the GFP control, (Fig. 6) indicates that CPR1 functions in early stages of infection. We also showed that soybean roots expressing CPR1 have enhanced susceptibility to SCN infection (Fig. 7B), further indicating a role in virulence. Surprisingly, roots expressing CPR1^C323S^ resulted in comparable cyst counts and cysts per gram of root to wild-type CPR1 expressing plants. We propose that CPR1^C323S^ is still able to associate with GmBCAT1, which is sufficient for disrupting function.

We initiated this work with the goal of generating PBS1-based decoy substrates for SCN effector proteases. We are currently working on identification of cleavage sequences preferred by CPR1. In the context of the PBS1 decoy system, however, it is notable that CPR1 can suppress ETI, which raises the question of whether resistance induced by cleavage of a GmPBS1 decoy will also be suppressed. The answer to this question may depend on the relative amount and timing of CPR1 delivery by the nematode. We speculate that efficient suppression of ETI may require large amounts of CPR1, as obtained by overexpression in *N. benthamiana*, whereas triggering of ETI requires only very small amounts of effector. We are thus optimistic that PBS1 decoys engineered to be cleaved by CPR1 will be effective at killing syncytial cells and thereby conferring resistance to SCN. Activation of cell death during the penetration process and/or during syncytium formation should restrict nutrient availability to the nematode, and thus again confer resistance. Furthermore, if decoy engineering is successful in preventing SCN parasitism, it should be possible to extend this system to confer resistance to other plant-parasitic nematodes that secrete effector proteases during infection.

## MATERIALS AND METHODS

### Plant growth and maintenance

*A. N. benthamiana* seeds were planted directly on Sun Gro propagation mix (Sun Gro Horticulture, Agawam, MA) supplemented with Osmocote^®^ 14-14-14 slow-release fertilizer. Plants were maintained in a growth room with a temperature of 21 to 23°C, a relative humidity ranging from (30-70%), and a 16-h light and 8-h dark photoperiod. For the composite soybean plants, soybean seeds [*Glycine max* (L.) Merr.] cultivar Williams 82 were planted directly on Pro-mix soil in 6-inch pots supplemented with Osmocote 14-14-14 slow-release fertilizer. Plants were maintained in a growth chamber set to a temperature of 22°C to 24°C, a relative humidity of 60%, and a 16-h light and 8-h dark photoperiod with an average light intensity at pot level of 300 µ Einsteins m^-2^ s^-1^. For penetration assays, *Glycine max* seeds (cultivar Williams 82) were surface sterilized with 70% ethanol for 2 min and then with 50% bleach for 10 min, followed by three rinses in sterile water. Sterilized seeds were placed on wet filter paper with 10 mM morpholinoethanesulfonic acid (MES) buffer [pH 6.5] inside a Petri plate and incubated in a growth chamber at 26°C for 5 days.

### Growth of bacterial strains

*Agrobacterium tumefaciens* GV3101 and *Agrobacterium rhizogenes* K599 strains were grown on Luria-Bertani (LB) plates for 2 days at 28°C in an incubator. Liquid LB cultures were grown at 28°C overnight on a shaker. *Escherichia coli* Top10 cells were grown overnight at 37°C either on LB plates (incubator) or LB liquid (shaker). Antibiotics were used at the following concentrations kanamycin 50 µg/ml, carbenicillin 100 µg/ml, gentamycin 10 µg/ml, and spectinomycin 50 µg/ml for selection during growth. All overnight incubation periods were for approximately 12 to 16 h.

### Generation of plant expression constructs

The coding sequence of *CPR1* (lacking the signal peptide) was synthesized and cloned into a pUC57-Amp plasmid (GeneWiz). For cloning *CPR1* constructs used in cleavage assays and localization, the coding sequence was amplified from the pUC57-Amp plasmid with primers adding attB1 and attB4 sites for Gateway-cloning into pBSDONR(P1-P4) (Qi et al. 2012). The pBSDONR-*CPR1* construct was sequence verified (Eurofins) and used as a template for site-directed mutagenesis to mutagenize the catalytic cysteine residue (CPR1^C323S^) or amplify without the N-terminal I29 inhibitor domain. Site-directed mutagenesis was performed as described in (Edelheit et al. 2009). pBSDONR-*CPR1* and pBSDONR-*CPR1^C323S^* served as templates to amplify the “mature” CPR1 variants with or without protease activity respectively. In brief, the mature variant (CPR1(mat)) was generated by PCR amplifying the coding sequence beginning at Ala 71, which is the first amino acid after the predicted I29 inhibitor domain. The PCR product was amplified with primers that incorporated a start codon and attB1-attB4 sites for Gateway cloning. Using multisite Gateway LR cloning, pBSDONR constructs carrying the different CPR1 variants (*CPR1, CPR1^C323S^, CPR1(mat), or CPR1(mat)^C323S^*) were cloned into dexamethasone (DEX) inducible vectors pBAV154 with a C-terminal miniTurbo fusion or pTA7001 with a C-terminal mCherry or sYFP2 fusion. All C-terminal protein fusions were sourced from pBSDONR (P4r-P2) vectors that harbored the coding sequences for the fluorescent proteins (Qi et al. 2012). These constructs were then transformed into *A. tumefaciens* GV3101. Gateway LR cloning was also used to assemble pTA7001-*AvrPphB^C98S^:eGFP*, pTA7001-*GmBCAT1:sYFP*, and pTA7001-*PBS1:mCherry*.

For the constructs used in the composite soybean plants, a combination of Gateway and Golden Gate assembly was utilized to insert multiple transcription units into a single T-DNA. Any coding sequence containing *Bbs*I or *Bsa*I sites was first adapted for use in the Golden Gate system by using site directed mutagenesis to remove endogenous *Bbs*I and *Bsa*I sites. The coding sequence of the three different variants of CPR1 were first fused with the miniTurbo tag and cloned into pBAV154 via Gateway cloning for consistency with previous constructs, then amplified from pBAV154 with primers to add overhangs for Golden Gate assembly, and finally cloned into the Level1 pICH47732 plasmid (Addgene catalog # 48000) with a CaMV 35S short promoter (pICH51277; Addgene catalog # 50268), and a CaMV 35S terminator (pICH41414; Addgene catalog # 50337). The screenable marker RUBY (He et al. 2020) was amplified from pCAMBIA2300 (previously generated and gifted by Sebastian S. Cocioba, Binomica Labs) with primers for Golden Gate assembly into the Level 1 pICH47742 plasmid (Addgene catalog # 48001) with a double 35S promoter (pICH51277; Addgene catalog # 50268) and the same 35S terminator. Level 1 constructs were assembled using *BsaI*-HFv2 restriction enzyme and T4 DNA Ligase. Level 2 constructs were assembled into pICH89921 with the pICH47732- *protease:miniTurbo*, pICH47742-*RUBY*, and end linker (pICH41744; addgene catalog # 48017) using a *BbsI*-HF restriction enzyme and T4 DNA Ligase. The MoClo Plant Parts Kit was a gift from Nicola Patron (Addgene kit # 1000000047) (Engler et al. 2014). The MoClo Toolkit was a gift from Sylvestre Marillonnet (Addgene kit # 1000000044) (Weber et al. 2011; Werner et al. 2012)

To generate the constructs used in the co-immunoprecipitation and cleavage assays (pTA7002-*BCAT1^Δ1-74^:4xMYC*, pTA7002-*GmBCAT1:4xMYC*, and pTA7002-*CPR1(mat)^C323S^:eGFP*), we used NEBuilder HiFi DNA Assembly Master Mix (New England Biolabs). pTA7002 was linearized using *XhoI* and *SpeI* restriction enzymes (New England Biolabs). *GmBCAT1* was PCR amplified with terminal extensions complementary to the resulting pTA7002 ends. For the full-length *GmBCAT1*, the N-terminus was synthesized by TwistBio, PCR amplified, and assembled. Each tag was amplified from the MoClo Plant Parts Kit plasmids with extensions complementary to the gene of interest (*GmBCAT1 or CPR1(mat)^C323S^*) at the N-terminus or the vector at the C-terminus. Primers for all constructs described can be found in Supplementary Table S2.

### RNA extractions and cDNA synthesis

Soybean root and leaf tissue was harvested from 7 to 14-day old soybean plants. Hairy root tissue expressing YFP:miniTurbo+RUBY was harvested 4 to 5 weeks post infection. All tissue was flash frozen in liquid nitrogen at the time of harvesting. Tissue was ground to a fine powder in a chilled mortar with a pestle under liquid nitrogen. Approximately 0.5 grams of tissue was transferred to a 2 ml microcentrifuge tube for RNA extraction. 1.0 ml of TRIzol reagent (Thermo Scientific) was added to the plant tissue powder and vortexed for 30 s. Samples were incubated on a rocker for 10 min at room temperature. 200 µl chloroform was added to the sample, vortexed for 30 s, and incubated on a rocker for 5 min at room temperature. Samples were then centrifuged at 12,000xg, 4°C, for 15 min. Approximately 400 µl of the supernatant was transferred to a new tube and mixed with 400 µl of isopropanol. Samples were incubated at a minimum of 2 h, with a maximum of overnight, at −20°C to precipitate the RNA. Samples were then centrifuged at 12,000xg, 4°C, for 10 min to pellet the RNA. The supernatant was carefully removed. The RNA pellet was washed 3 times with 1 ml 75% ethanol, vortexed to mix and centrifuged at 7,500xg, 4°C, for 5 min. The pellet was dried for approximately 15 min under a sterile hood before resuspending in 50 µl of water. The RNA concentration was measured using a Nanodrop One instrument (Thermo Scientific). A DNase treatment was performed with 2 µg of the RNA prior to cDNA synthesis. 1 µg of RNA, the Thermo Verso cDNA Synthesis Kit (Thermo Scientific), and an Anchored Oligo dT primer was used to generate soybean cDNA. The resulting cDNA concentration was measured using the Nandrop One and used as a template for cloning of soybean genes.

### Transient expression in *N. benthamiana*

*A. tumefaciens* (GV3101) cells were scraped from Luria-Bertani (LB) plates and resuspended in 10 mM MgCl_2_. Final OD_600_ were adjusted based upon experiment and described under specific method. Resuspensions were supplemented with 100 µM acetosyringone, gently mixed by inverting, and incubated at room temperature for 2 to 3 h. After incubation, cultures were infiltrated into leaves of 4 to 6-week-old *N. benthamiana* plants with a needleless syringe. Plants were sprayed with 50 µM dexamethasone (Thermo Scientific), 0.02% Tween-20, 40 to 45 h post infiltration and tissue were harvested for analysis 4 to 21 h post induction, as specified in figure legends.

### Cell death suppression and electrolyte leakage assay

Mixtures of *A. tumefaciens* carrying pTA7001-RPS5^D266E^:5xMYC (OD_600_ of 0.150) and pTA7001-CPR1 variants with a C-terminal mCherry tag (OD_600_ of 0.400) were prepared for transient expression. Gene expression was induced 40 to 45 h post infiltration via dexamethasone. For visual analysis of cell death, images were taken under white and ultraviolet light 24 h post gene expression induction. For each replicate, ten to twelve leaves were injected from at least six different plants. Plants exhibiting tissue collapse prior to gene expression induction were not scored or imaged for cell death. At least two independent replicates were performed for all cell death assays.

For electrolyte leakage, 3 leaf discs per treatment from 4 different leaves were rinsed three times in deionized water approximately 3.5 h post gene expression induction. Leaf discs were floated in sterile deionized water with 0.001% Tween-20. Measurements were taken with a conductivity meter (HORIBA Scientific) at 4, 6, 8, 12.5, 15, and 26 h post gene expression induction. Measurements for each condition and timepoint were imported into GraphPad to calculate the standard error of the mean (SEM) and generate a graph.

### Protein isolation and immunoblots

For protein isolation from *N. benthamiana,* leaf tissue was harvested 4 to 21 h post gene expression induction via dexamethasone, weighed, and flash frozen in liquid nitrogen. Tissue was ground in ice-cold protein extraction buffer (150 mM NaCl, 50 mM Tris-HCl [pH 7.5], 0.1% Nonidet P-40 [Sigma-Aldrich], 1% protease inhibitor cocktail [Sigma-Aldrich], 2 mM 2,2’-dipyridal disulfide) with a chilled mortar and pestle. Cell debris was removed by centrifuging twice at 10,000xg, 4°C, for 10 min. *N. benthamiana* proteins were prepared in sodium dodecyl sulfate (SDS) loading buffer supplemented with 5% beta-mercaptoethanol and heat denatured in a heating block at 95°C for 10 min. The proteins were separated on a 4 to 20% Trist-glycine stain free polyacrylamide 10-well gel (Bio-Rad) at 170 V for approximately 1 h in 1x Tris/glycine/SDS running buffer. Loading controls were obtained by imaging the stain-free polyacrylamide gel using the stain-free gel setting on a ChemiDoc^TM^ Imaging System (Bio-Rad). Proteins were transferred to nitrocellulose membrane (Cytiva) at 300 milliamps for 1 h. Membranes were stained in Ponceau for approximately 1 min, washed in deionized water, and imaged.

Membranes were blocked in 5% nonfat dry milk (w/v) in 1x TBS-T overnight at 4°C on a shaker. Membranes were then incubated for 1 h at room temperature on a rocker with horseradish peroxidase (HRP)-conjugated c-MYC antibody (Invitrogen; Cat. No. MA1-81357), (HRP)-conjugated HA antibody (Roche; Cat. No. 12013819001), GFP monoclonal antibody (Proteintech; Cat. No. 66002-1-Ig), or RFP monoclonal antibody (Chromotek; Cat. No. 6G6) to detect proteins. Membranes were washed three times for 15, 5, and 5 min in 1x Tris-buffered saline (TBS) with 0.1% Tween-20 (TBS-T). The anti-GFP blots and anti-RFP blots were then incubated with HRP-conjugated goat anti-mouse antibody (Abcam) for 1 h at room temperature on a rocker. The anti-MYC, anti-RFP, and anti-mouse antibodies were diluted to a concentration of 1:5000 in 5% milk. The anti-GFP antibody was diluted to a concentration of 1:1000 in 5% milk. Membranes were washed an additional three times in 1x TBS-T after which they were incubated with ProtoGlow chemiluminescent substrate (National Diagnostics) for 5 min and exposed on ChemiDoc^TM^ Imaging System (Bio-Rad) using custom settings for exposure time.

### Fluorescence microscopy

Proteins tagged with sYFP or mCherry were transiently expressed in *N. benthamiana* using a dexamethasone inducible promoter. 40 to 45 h post *A. tumefaciens* infiltration, attached leaves were sprayed with 50 µM dexamethasone and 0.01% Tween-20. At 5 to 8 h post transgene induction, leaves were imaged using a Leica Stellaris 8 FALCON Confocal microscope with a 63x water objective. For sYFP (when co-expressed with mCherry), a 514 nm excitation wavelength was used, and the images were viewed between 526 to 569 nm wavelength of emission. For mCherry, a 587 nm excitation wavelength was used, and the images were viewed between 600 to 640 nm wavelength of emission. For MitoTracker Red (Molecular Probes), lyophilized powder was resuspended in dimethyl sulfoxide to a concentration of 1 mM. A working solution of 100 µM was prepared in 10mM MgCl_2_ and infiltrated into *N. benthamiana* leaves 4 to 6 h post transgene induction and at least 30 min prior to imaging. Leaves were incubated and kept in the dark. For MitoTracker Red, a 579 nm excitation wavelength was used, and the images were viewed between 590 to 620 nm wavelength of emission. When sYFP was co-expressed with MitoTracker Red a 513 nm excitation wavelength was used and the images were viewed between 523 to 540 nm wavelength of emission. A minimum of three leaves were imaged between at least two independent replicates.

### Generation of composite soybean plants

The protocol for generation of composite plants was optimized from (Fan et al. 2020), with minor modifications. In brief, *A. rhizogenes* K599 strains carrying T-DNA constructs harboring protease variants and the RUBY screenable marker were prepared on LB solid and in LB liquid media. Prior to infection, liquid cultures were centrifuged at 4,000xg for 5 min, the pellet was washed once with ¼-strength Gamborg’s liquid [pH 5.75] (plantMedia), and re-centrifuged. Bacterial pellets were resuspended in ¼ strength Gamborg’s [pH 5.75] (plantMedia) and adjusted to an OD_600_ of 1.000 to 1.200. Slant cuts 0.5-1 inch long were made on hypocotyls of 7-day old soybean seedlings, right below the cotyledons, with a sterile razor. *A. rhizogenes* K599 strains were scraped from plates and lathered on the cut site of the scion. The scions were then placed in 1020 plastic flats filled with coarse vermiculite soaked in ¼-strength Gamborg’s media [pH 5.75]. The infected plants were maintained under clear plastic humidity bags to maintain high humidity inside a growth chamber set to 23 to 24°C, 60% humidity, and a 16-h light/8-h dark photoperiod for the first 7 to 9 days. Plants were carefully removed from the vermiculite at approximately 10 days post inoculation *with A.* rhizogenes K599 strains to remove non-transgenic (non-red) tissue. Plants were maintained under standard soybean growth conditions (described above) in vermiculite or soil from 3 weeks post infection.

### Proximity-based labeling in composite soybean plants

The protocol for miniTurbo-mediated proximity labeling was optimized from (Zhang et al. 2020) for use in soybean roots. In brief, transgenic soybean roots expressing the visual RUBY marker were submerged in 250 µM biotin (prepared in dimethyl sulfoxide) in ¼-strength Gamborg’s. The roots were vacuum infiltrated for 1 min and maintained in a growth chamber for 4 h. Beakers were covered in foil to prevent light exposure to roots during the incubation. Approximately 1.5-2.0 gram of tissue was harvested from each sample set and flash frozen in liquid nitrogen. Proteins from root tissue were extracted in two volumes of RIPA buffer (50 mM Tris-HCl [pH 7.5], 500 mM NaCl, 1 mM EDTA, 1% NP40 [v/v], 0.1% SDS [w/v], 0.5% sodium deoxycholate [w/v], 1 mM DTT, 1% protease inhibitor cocktail [Sigma Aldrich]). Free biotin was removed from the samples using 10 ml Zeba^TM^ Spin Desalting Columns 7K MWCO (Thermo Scientific). Bradford analysis was used to quantify protein concentration post desalting. Samples were adjusted to 4 mg/ml and then incubated with 100 µl of streptavidin Dynabeads overnight at 4°C. The beads were collected on a magnetic rack for 5 min and the protein supernatant was removed. A series of washes were performed as described in (Zhang et al. 2020). The beads were captured on a magnetic rack for 5 min between each wash. All washes were performed at room temperature. The last wash buffer (50 mM ammonium bicarbonate) was removed, and the beads were frozen at −80°C until proteins were digested and analyzed via mass spectrometry.

### Protein digestion and mass spectrometry

Individual samples containing streptavidin beads and associated proteins were denatured in 8 M urea in 100 mM ammonium bicarbonate. Samples were incubated for 45 min at 57°C with 10 mM Tris(2-carboxyethyl)phosphine hydrochloride to reduce cysteine residue side chains. These side chains were then alkylated with 20 mM iodoacetamide for 1 h the dark at 21°C. The urea was diluted to 1 M using 100 mM ammonium bicarbonate. A total of 0.4 µg trypsin (Promega) was added, and the samples were digested for 14 h at 37°C.

The resulting peptide solution was centrifuged at 2000 rcf for 1 min to pellet the beads. The supernatant was transferred to fresh tubes and desalted using ZipTip pipette tips (EMD Millipore), dried down and resuspended in 0.1% formic acid. Peptides were analyzed by LC-MS on an Orbitrap Fusion Lumos equipped with an Easy NanoLC1200. Buffer A was 0.1% formic acid in water. Buffer B was 0.1% formic acid in 80% acetonitrile. Peptides were separated on a 90-min gradient from 0% B to 35% B. Peptides were fragmented by HCD at a relative collision energy of 32%. Precursor ions were measured in the Orbitrap with a resolution of 60,000. Fragment ions were measured in the Orbitrap with a resolution of 15,000.

Data were analyzed using Proteome Discoverer (2.5) to interpret and quantify the relative amounts in a label free quantification manner. Data was searched against the *Glycine max* proteome downloaded on 10/26/2022. Trypsin was set as the protease with up to two missed cleavages allowed. Carbamidomethylation of cysteine residues was set as a fixed modification. Oxidation of methionine and protein N-terminal acetylation were set as variable modifications. A precursor mass tolerance of 10 ppm and a fragment ion quantification tolerance of 0.05 Da were used. Data was quantified using the Minora feature detector node within Proteome Discoverer.

### Co-immunoprecipitation and cleavage assays

For co-immunoprecipitations (co-IPs), *N. benthamiana* leaves co-expressing the bait (CPR1(mat)^C323S^:GFP or AvrPphB^C98S^:GFP) and the putative soybean target protein, with C-terminal 4xMYC protein fusions, were harvested 21 h post gene expression induction for protein isolation. Bait proteins were infiltrated at an OD_600_ of 0.400 and 0.300 respectively. GmBCAT^1-74^ was infiltrated at an OD_600_ of 0.500. Protein was extracted in ice-cold co-IP extraction buffer (10% glycerol, 25 mM Tris-HCl [pH 7.5], 1 mM EDTA, 150 mM NaCl, 10 mM dithiothreitol, 1x protease inhibitor cocktail [Sigma-Aldrich], 1 mM phenylmethylsulfonyl fluoride), 1 ml of protein lysate was incubated with 20 µl GFP-trap agarose (Chromotek) beads that were previously equilibrated with co-IP extraction buffer according to manufacturer’s directions at 4°C, on a rotator, for 2 h. Beads were washed 5 times with co-IP extraction buffer supplemented with 0.1% NP-40. After the third wash, beads were transferred to a new 1.5 ml microcentrifuge tube for the final two washes. For analysis of proteins captured on GFP-trap beads for co-IP assays, beads were boiled in SDS loading buffer + 10% beta-mercaptoethanol at 95°C for 10 min. Samples were centrifuged at 2,500xg for 3 min to separate beads and protein samples and then analyzed by immunoblot analysis as described above. The co-IP assays were repeated three times.

For cleavage assays, *N. benthamiana* leaves co-expressing the CPR1 variants with a C-terminal mCherry fusion protein (OD= 0.400) and GmBCAT1 (or GmBCAT1^Δ1-74^) with a C-terminal sYFP or 4xMYC protein fusion (OD= 0.500), were harvested at 4 and/or 21 h post gene induction for immunoblot analysis. Cleavage tests were performed at least two times.

### Phylogenetic analysis of *Arabidopsis* and soybean BCAT proteins

The Arabidopsis BCAT1 amino acid sequence (At1g10060) was downloaded from the NCBI Protein database (Genbank accession number BAH19488). The proteomes of soybean (*Glycine max* W82.a6.v1, Valliyodan et al. 2019) and Arabidopsis (Araport11, Cheng et al. 2017) were obtained from the Phytozome 13 online database (Goodstein et al. 2012). These were then queried with the Arabidopsis BCAT1 sequence using BLASTp (2.13.0+) (Altschul et al. 1990; Altschul et al. 1997). We retained all proteins with alignment scores above 300. Secondary protein isoforms were removed, retaining the best scoring isoform from each gene. Amino acid alignments were created using Clustal Omega 1.2.4 (Sievers and Higgins 2021). A Maximum Likelihood tree was created using RaxML 8.2.12 in PROTGAMMAAUTO mode with 1000 bootstraps (Kozlov et al. 2019). Trees were displayed usig iTOL v6.9.1 (Letunic and Bork 2024), with BCAT-like genes pruned from the tree due to more distant evolutionary relationships.

### dsRNA synthesis and purification

A 289 bp fragment of the *CPR1* gene and a 255 bp dsRNA targeting a synthetic GFP (Green Fluorescent Protein) gene were amplified using primers with T7 promoter sequences added to their 5’ ends (Supplementary Table S3). These PCR products were then used for in vitro transcription using T7 RNA polymerase. The synthetic GFP gene, not naturally present in the nematode, served as a negative control in the experiments. To facilitate octopamine-induced ingestion by nematodes, the size of dsRNA for both genes was kept below 300 bp.

Sense and antisense RNA were synthesized in a single in vitro reaction using the MEGAscript® RNAi Kit (Thermo Scientific) according to the manufacturer’s instructions, with an incubation period of 8 h to enhance RNA yield. The resulting dsRNA product was purified using an ethanol precipitation protocol (Green and Sambrook 2020) and its integrity assessed using agarose gel electrophoresis. Its concentration and quality were assessed using a Nanodrop spectrophotometer (Thermo Scientific).

### RNA interference

RNAi soaking was conducted following the protocols outlined by (Sukno et al. 2007) with slight modifications. Approximately 10000 nematodes were used per biological replicate of freshly hatched *H. glycines* J2 worms, soaked in a mixed buffer containing 2 mg/ml and 3 mg/ml dsRNA in 1/4 M9 buffer (43.6 mM Na_2_HPO_4_, 22 mM KH_2_PO_4_, 2.1 mM NaCl, 4.7 mM NH_4_Cl), 1 mM spermidine, and 50 mM octopamine at 26°C on a rotator covered with aluminum foil to maintain a dark environment. J2 worms incubated in dsRNA of the GFP gene were utilized as the control. After 24 h of incubation, J2 worms were washed three times with Nemawash (10 mM MES buffered water [pH 6.5], 0.01% Tween-20) through brief centrifugation to remove external dsRNA. Three biological replicates were used for each gene.

### Reverse transcription quantitative PCR (RT-qPCR)

RT-qPCR was employed to investigate the impact of RNAi on the expression of CPR1. Total RNA was isolated from *H. glycines* J2 worms following treatment with CPR1-dsRNA or GFP-dsRNA control using the Nucleospin microRNA kit (Macherey-Nagel, Hoerdt, France). First-strand cDNA was synthesized and served as a template for PCR. RT-qPCR was conducted using iTaq universal SYBR Green super mix (Bio Rad) on a Bio Rad CFX96 Real-Time PCR Machine. Each reaction included 2 µl of each primer [10 pM/µl], 5 µl of cDNA, 10 µl SYBR Green super mix, and 1 µl of nuclease-free water, making a final volume of 20 µl. Thermocycler conditions comprised an initial denaturation cycle at 94°C for 30 s, followed by 40 cycles at 94°C for 5 s and 58°C for 34 s. The reaction concluded with the determination of the dissociation curve for all amplicons. The experimental design incorporated three biological replicates and three technical replicates. GAPDH and Actin served as the internal control to normalize the reaction (Supplementary Table S3).

### Nematode penetration assays

Five-day-old surface sterilized soybean seedlings were used for the experiments. A 23% Pluronic F-127 (PF-127) (Sigma-Aldrich) gel was prepared. SCN infection was assessed in a 6-well tissue culture plate. Four milliliters of Pluronic gel were poured into each well, and seedlings were placed in each well at 15-20°C. After the gel solidified, approximately 120 J2s/50 µl of *H. glycines* (TN10) were inoculated at the root tip of each seedling using a pipette tip, and plates were stored at 28°C for 24 h. Three biological replicates were utilized for each treatment, totaling 47 plants for GFP and 47 plants for CPR1 in the analysis. Eight plates were employed for each treatment in the experiment. After soaking, nematodes were washed twice with Nemawash solution (10 mM MES buffer [pH 6.5], 0.01% Tween-20). Subsequently, the nematodes were counted four times, and the final average of the four replicates were calculated. The concentration of the nematodes was adjusted to 120 J2 suspended in 50 μl of Pluronic gel (120 J2s/50 μl). After 24 h, plants were harvested from the gel by briefly positioning the plates over an ice bath. This slight decrease in temperature caused the gel to liquify, facilitating the extraction of plantlets without causing damage to the root system. For staining, roots underwent a 2-min treatment with 1% bleach followed by immersion in boiling acid fuchsin solution (Sigma-Aldrich) for 2 to 3 min. Subsequently, the roots were immersed in acidified glycerol (10 drops) and left for destaining. Nematodes were counted under a dissecting microscope.

### Nematode infection assays of composite soybean plants

Transgenic composite soybean plants were removed from vermiculite again at approximately 21 days post inoculation with *A. rhizogenes* K599 strains to remove non-transgenic tissue as before, then were placed in 8-inch conetainers filled with a mixture of two-parts sand to one-part field soil that had previously been steam sterilized. Plants were allowed to recover for two additional days and then inoculated with 2000 eggs of SCN population TN10. Nematodes were allowed to infect and develop to adulthood (approximately 28 days), at which time the root systems were removed from conetainers and the cysts from each root system were collected in a manner similar to (Kandoth et al. 2011). Wet weight measurements were recorded for each root system. Cysts from replications of each transgenic construct were transferred to individual wells of 12 well plates. Each well was then photographed using a Zeiss Stemi SV11 microscope with AxioCam HRc and Axiovision SE64 V4.9.1 software at 1040x1040 pixel resolution and each photograph (see representative photograph) was used as input for an AI software-based cyst counting program (Nemacounter) developed in house (https://github.com/DjampaKozlowski/NemaCounter). Cysts per gram wet weight were generated for replications of each construct.

## ACKNOWLEDGEMENTS

The authors would like to thank undergraduates O. Yeley (Indiana University) and J. Goshon (Iowa State University) for assisting in data collection and plant maintenance; S. Ghosh for guidance and training with confocal microscopy; S. Pottinger and A. Sartor Chicowski for input on miniTurbo optimization; S. Cocioba for gifting the pCAMBIA-RUBY plasmid that was used as a template for our cloning; S. Eves-van den Akker for engaging discussions; the Indiana University Light Microscopy Imaging Center and A. Kun for assistance with light microscopy; the Indiana University Biological Mass Spectrometry Facility and J. Trinidad for providing mass-spectrometry based analyses; and the Indiana University Greenhouse and J. Leichter for assistance with growing plants.

## Supplementary Figures

**Supplementary Fig. S1.**
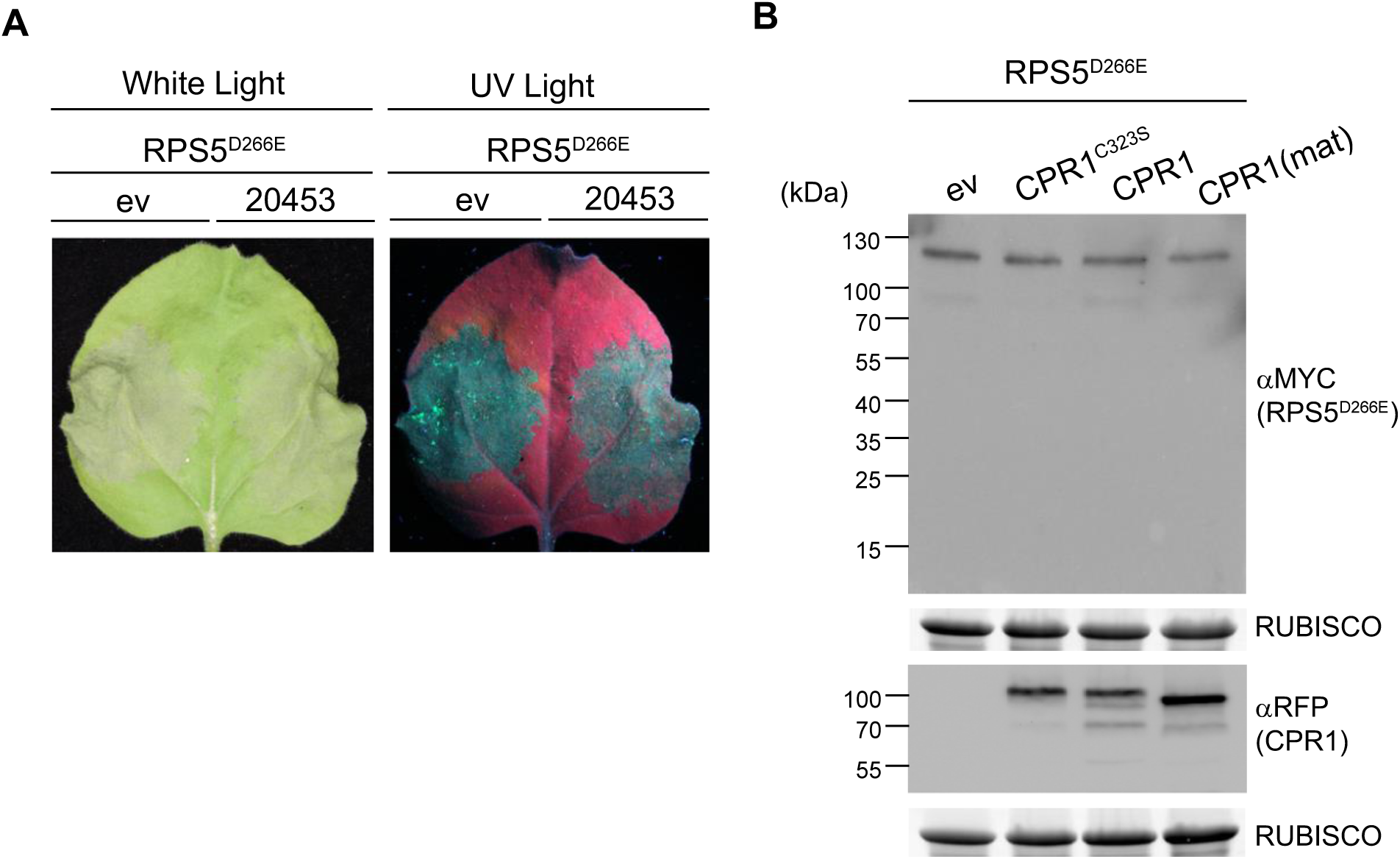
**A,** Screening SCN protease (20453) for HR suppression using the RPS5^D266E^ assay. 20453 was screened for HR suppression in parallel to CPR1 (shown in Fig. 1A) and was unable to suppress HR. 10/10 leaves resulted in tissue collapse in both the empty vector (ev) control and the infiltration area expressing 20453:mCherry. Images were taken 24 h post gene expression induction. **B,** The Immunoblot analysis of representative tissue from electrolyte leakage assay. Immunoblot of RPS5^D266E^:5xMyc and CPR1:mCherry variants confirms expression of proteins transiently co-expressed in *N. benthamiana* leaves. Samples were harvested in parallel to the first electrolyte leakage reading; approximately 4 h post gene expression induction. RUBISCO was used as a loading control.

**Supplementary Fig. S2.**
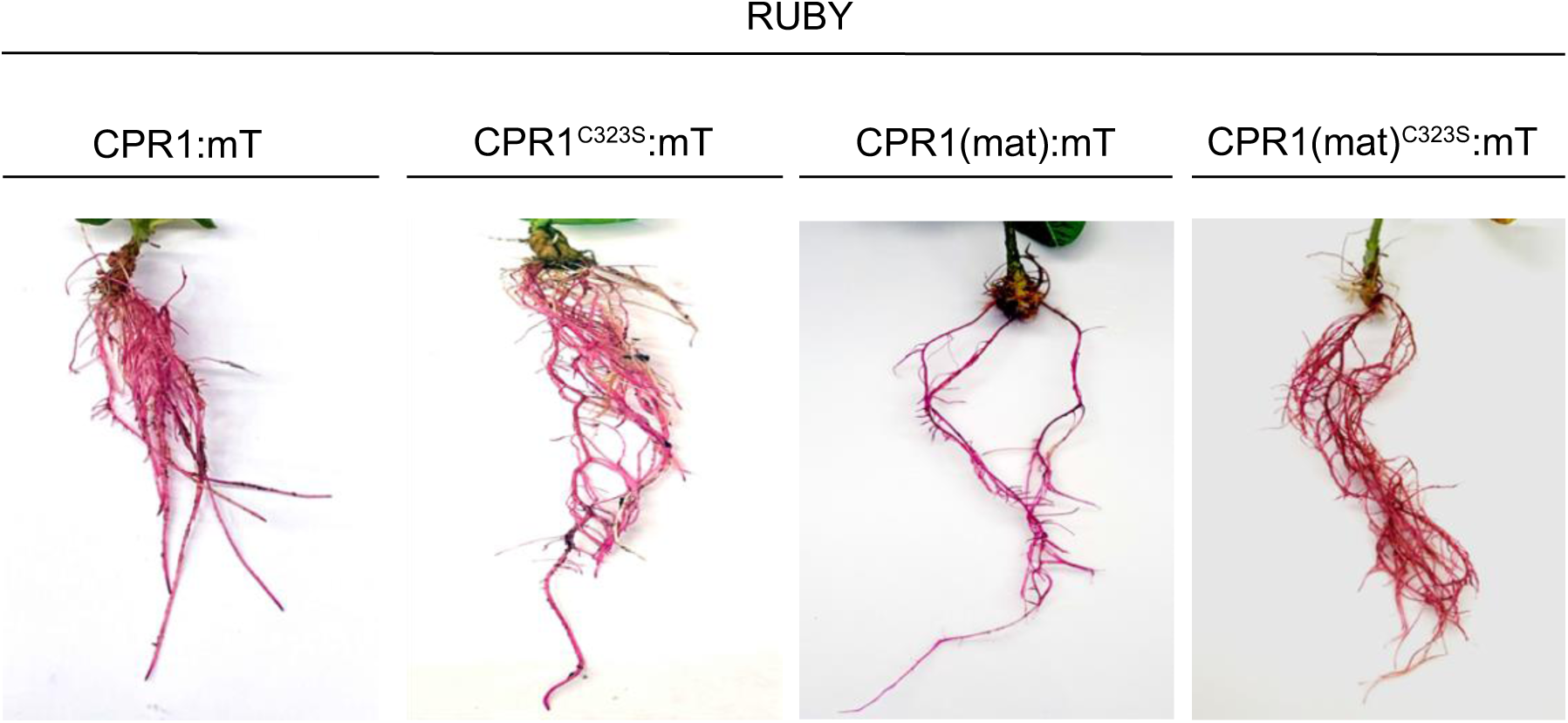
Composite soybean plants expressing CPR1:miniTurbo + RUBY variants for proximity-based labeling experiments. Images of composite soybean roots expressing CPR1:miniTurbo+RUBY variants. Images were taken 4-5 weeks post infection for hairy root induction. Images are representative of 12 plants generated for each construct. Roots were generated at least three times for each CPR1 variant with consistent results.

**Supplementary Fig. S3.**
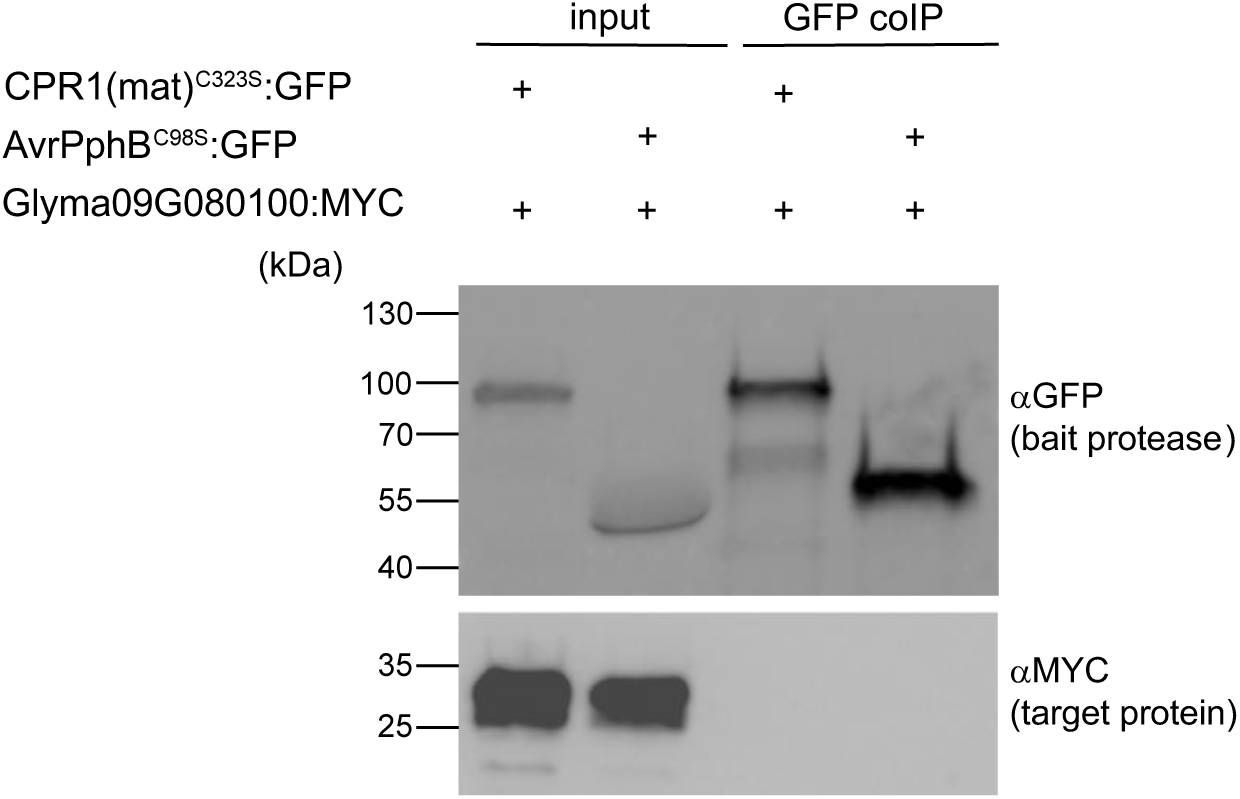
CPR1 does not co-immunoprecipitate with Glyma09G080100. This co-IP was performed in parallel to the co-IP shown in Fig. 4A, which showed that CPR1 co-IPs with GmBCAT.

**Supplementary Table S1.** Top priority targets identified using miniTurbo-based proximity labeling.

| UniprotID | Gene Name | Annotation |
| --- | --- | --- |
| K7KT64* | GLYMA_06G050100 | Branched-chain-amino-acid aminotransferase |
| A0A0R4J3C5* | GLYMA_05G182600 | Alanine--glyoxylate aminotransferase 2 homolog 1, mitochondrial |
| A0A0R0IBM9* | GLYMA_09G080100 | CMP/dCMP-type deaminase domain-containing protein |
| I1LMW7* | GLYMA_11G234500 | Alpha-soluble NSF attachment protein (SNAP11) |
| I1JZK8 | GLYMA_05G023100 | t-SNARE coiled-coil homology domain-containing protein (SNAP25) |
| C6T588 | GLYMA_07G243600 | Bet_v_1 domain-containing protein |
| I1KVZ9 | GLYMA_08G230100 | Bet_v_1 domain-containing protein |
| I1JJM7* | GLYMA_02G303300 | galactinol--sucrose galactosyltransferase |
| A0A0R4J2Q8* | GLYMA_02G217100 | Serine hydroxymethyltransferase |
| I1JE17 | GLYMA_02G104600 | Glycosyltransferase |
| C6SWE6* | GLYMA_10G151800 | TCTP domain-containing protein |
| A0A0R0H3D0 | GLYMA_12G092900 | Secoisolariciresinol dehydrogenase-like |

\* genes that were successfully cloned from soybean root cDNA.

**Supplementary Table S2.**
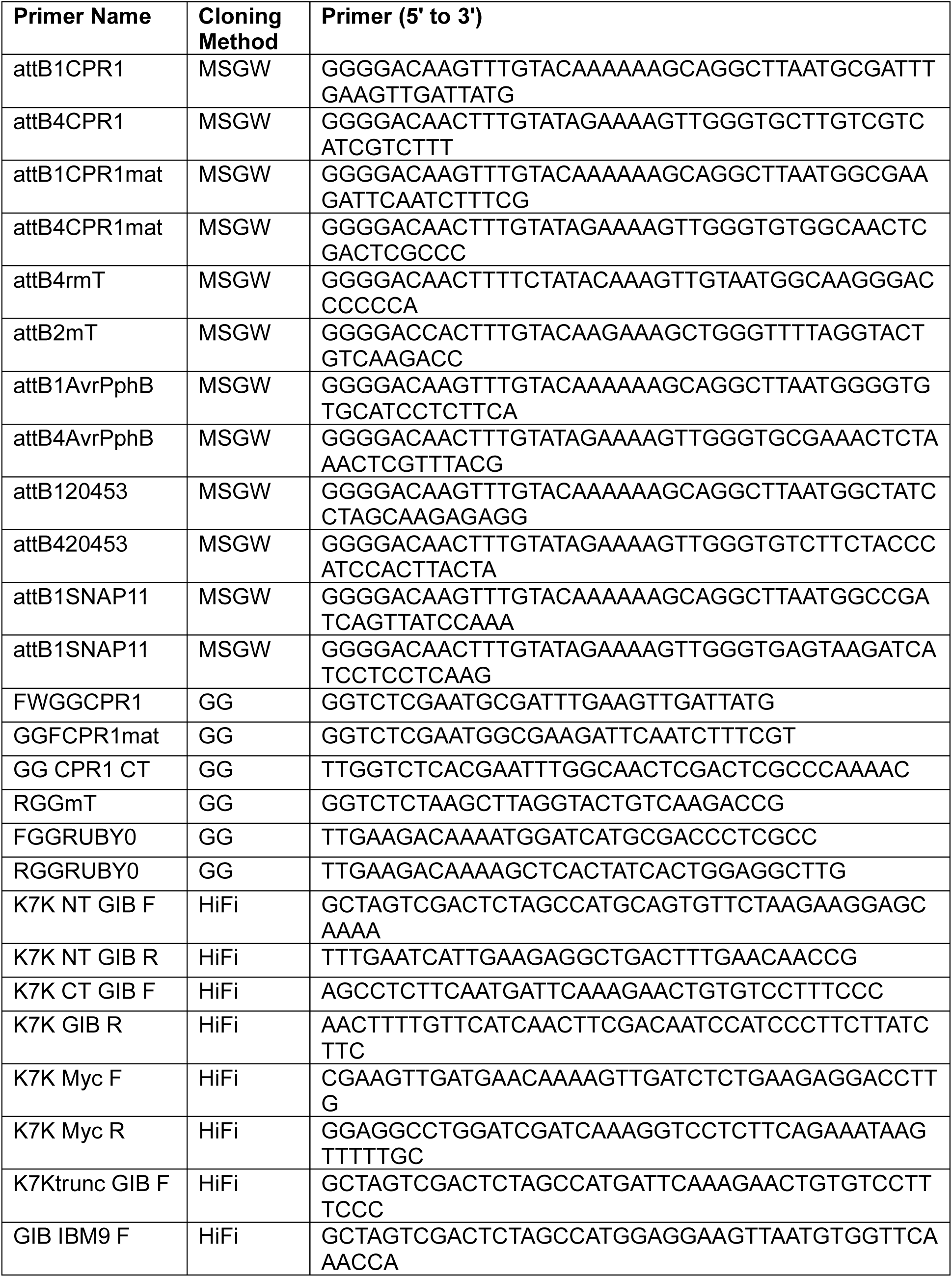
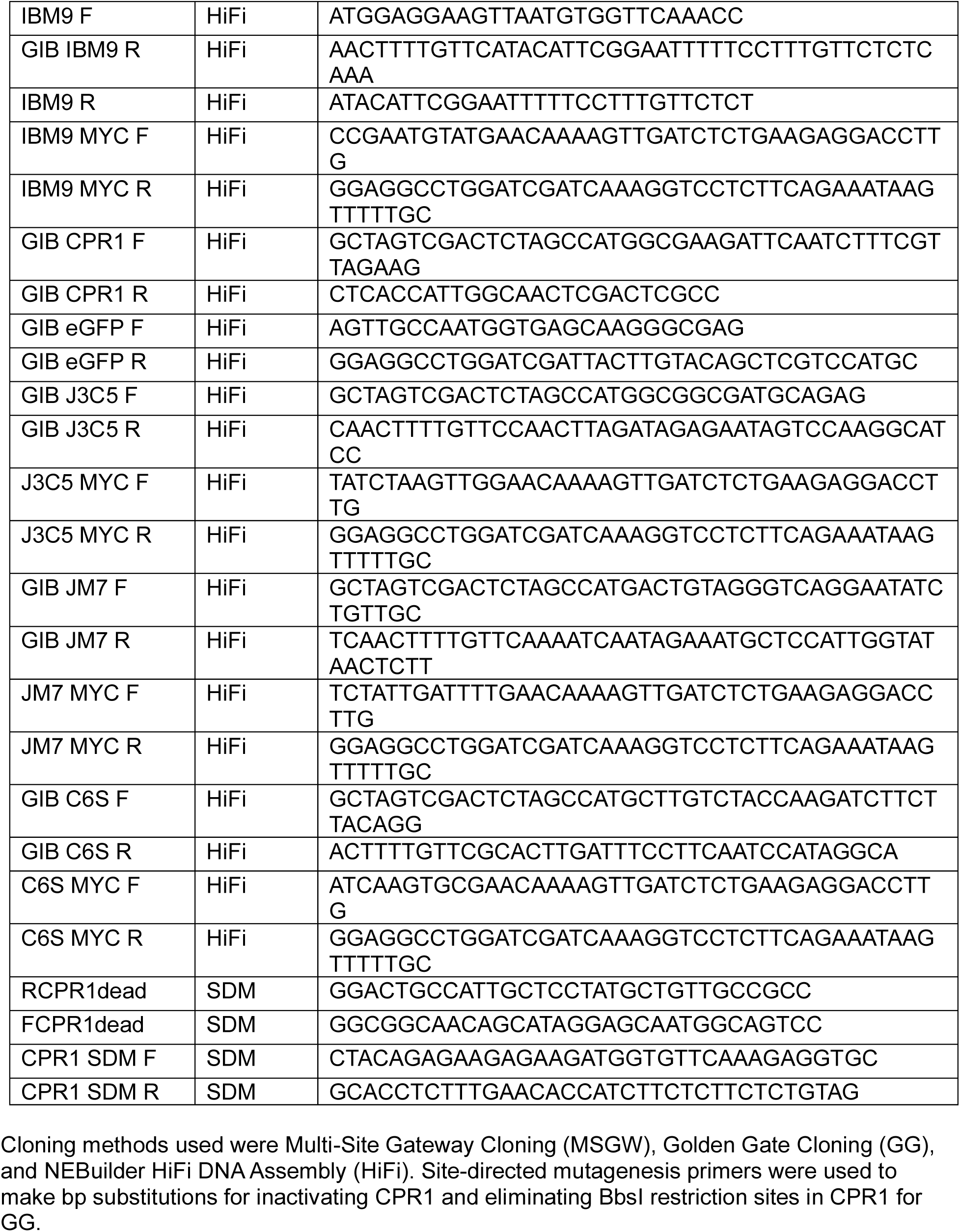
Primers used for generation of constructs.

**Supplementary Table S3.**
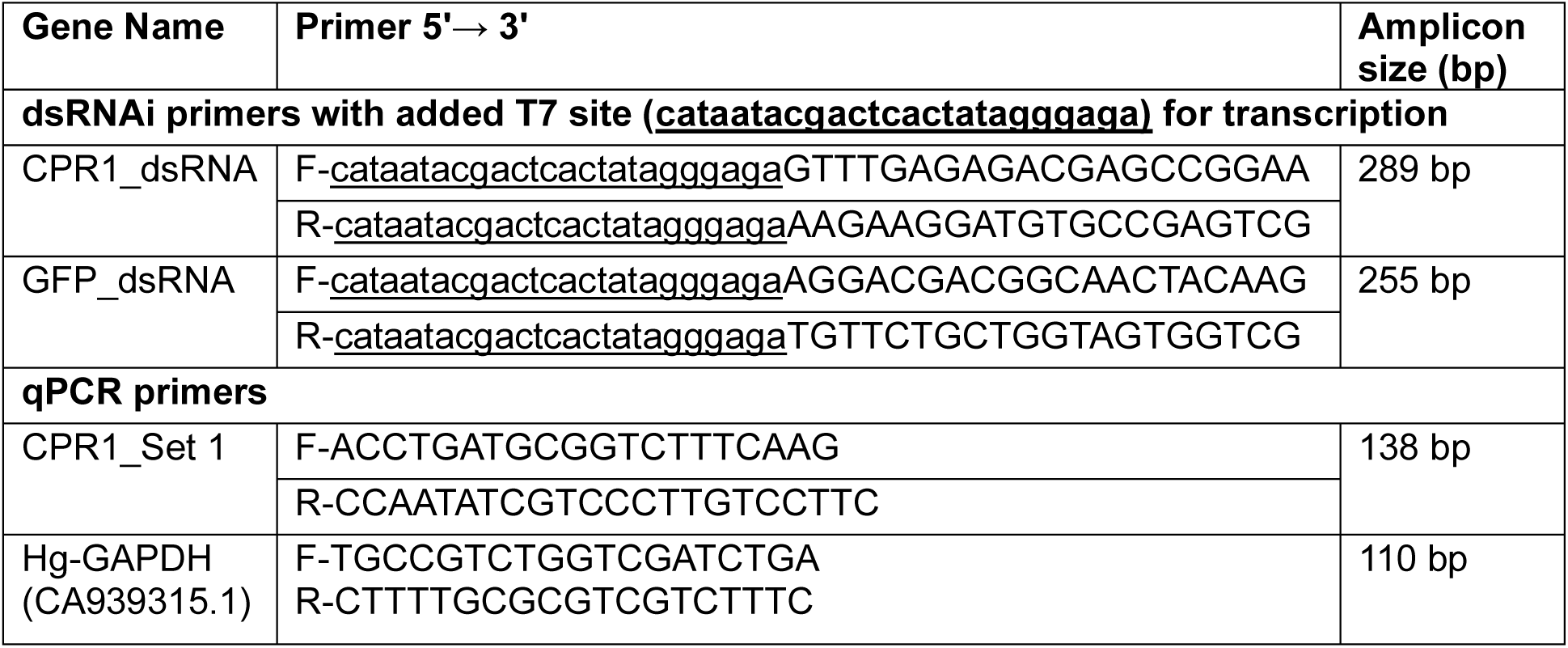
Primers used for gene silencing and qPCR.

